# Updated knowledge and a proposed nomenclature for nuclear receptors with two DNA Binding Domains (2DBD-NRs)

**DOI:** 10.1101/2023.05.09.540016

**Authors:** Wenjie Wu, Philip T. LoVerde

## Abstract

Nuclear receptors (NRs) are important transcriptional modulators in metazoans. Typical NRs possess a conserved DNA binding domain (DBD) and a ligand binding domain (LBD). Since we discovered a type of novel NRs each of them has two DBDs and single LBD (2DBD-NRs) more than decade ago, there has been very few studies about 2DBD-NRs. Recently, 2DBD-NRs have been only reported in Platyhelminths and Mollusca and are thought to be specific NRs to lophotrochozoan. In this study, we searched different databases and identified 2DBD-NRs in different animals from both protostomes and deuterostomes. Phylogenetic analysis shows that at least two ancient 2DBD-NR genes were present in the urbilaterian, a common ancestor of protostomes and deuterostomes. 2DBD-NRs underwent gene duplication and loss after the split of different animal phyla, most of them in a certain animal phylum as paralogues, rather than orthologues, of that in another animal phylum. Amino acid sequence analysis shows that the conserved motifs in typical NRs are also present in 2DBD-NRs and they are gene specific. From our phylogenetic analysis of 2DBD-NRs and following the rule of Nomenclature System for the Nuclear Receptors, a nomenclature for 2DBD-NRs is proposed.

## INTRODUCTION

Nuclear receptors (NRs) are important transcriptional modulators in metazoans, members of the NR superfamily are characterized by a modular structure: typical NRs contain an N terminal A/B domain, a C domain (DNA binding domain, DBD), a D domain (hinge) and an E domain (ligand binding domain, LBD) (Fig. 1A). NRs regulate transcription through binding to the promoter region of their target gene by the DBD and activation or repression of mRNA synthesis through co-regulators bound to the LBD [1–5]. Atypical NRs exist in some animals, NRs with DBD but no LBD are found in arthropods and nematodes, NR missing DBD but contain a LBD are present in vertebrates (Fig. 1A).

**Figure 1.**
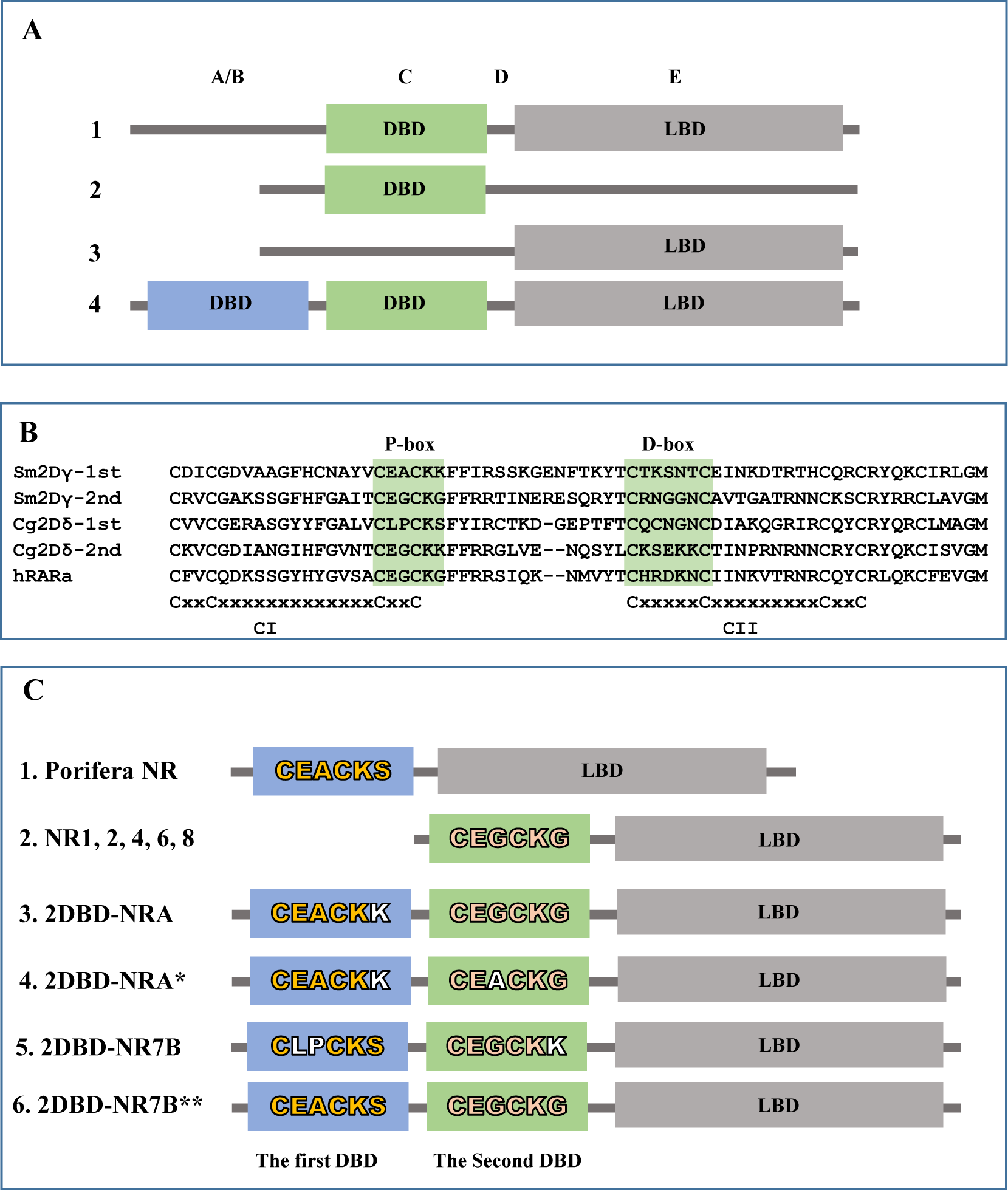
Modular structure of nuclear receptor. **A.** Modular structure of nuclear receptors. 1. Typical NR with single DBD and a LBD, 2. Atypical NR with only a DBD but without LBD, 3. Atypical NR with only a LBD but without DBD, 4. Atypical NR with two DBDs and a LBD. **B**. DBD sequence alignment shows the two zinc fingers and the conserved P-box and D-Box. The first zinc finger (CI) with a conserved motif sequence of C-X2-CX13-C-X2-C; the second zinc finger (CII) with a conserved motif sequence of C-X5-C-X9-C-X2-C. C: cysteine residue, X followed by a number that indicates the number of amino acids between the Cs (Cys). Sm2Dγ-1st: the first DBD of *Schistosoma mansoni* 2DBD-NRγ (Sm2DBD-NRγ, GenBank: AAW88550), Sm2Dγ-2nd: the second DBD of Sm2DBD-NRγ, Cg2Dδ-1st: the first DBD of *Crassostrea gigas* 2DBD-NRδ (Cg2DBD-NRδ, GenBank: XP_011428801), Cg2Dδ-2nd: the second DBD of Cg2DBD-NRδ, hRARα: human RARα (GenBank: AAD05222.1). **C**. P-P module of 2DBD-NRs (the P-box sequence in the first DBD and the second DBD of 2DBD-NRs). 1. Shows the P-box sequence of a Porifera NRs (*Suberites domuncula* RXR, SdRXR, GenBank: CAD57002.1) that is same as the first DBD of Echinodermata 2DBD-NRB. 2. Shows the P-box sequence of NRs in subfamily 1, 2 4, 6 and 8 that is same as the P-box of the second DBD of 2DBD-BRA. 3. P-P module of 2DBD-NRA (CEACKK-CEGCKG) which is found in most of 2DBD-NRA except members of Rotifers NR7A3 and NR7A5 groups. 4. P-P module of 2DBD-NRA (CEACKK-CEACKG) which is only found in the members of Rotifers NR7A3 and NR7A5 groups. 5. P-P module of 2DBD-NRB (CLPCKS-CEGCKK) which is found in most of 2DBD-NRB except members of Echinodermata 2DBD-NRB. 6. P-P module of 2DBD-NRB (CEACKS-CEGCKG) which is only found in Echinodermata 2DBD-NRB.

In 2006, we reported our result of identification and isolation of partial cDNAs of three NRs from blood fluke *Schistosoma mansoni*, each of them possesses two tandem DBDs (2DBD-NRs) [6], they were then verified by the *S. mansoni* Genome Project [7] and the full length cDNAs were isolated [8]. This was the first time to demonstrate that NR possesses a novel modular structure: A/B-DBD-DBD-hinge-LBD organization [8] (Fig. 1A). By an extensive search of whole genomic sequence (WGS) databases, we further demonstrated 2DBD-NRs were present in other animals including Platyhelminths *Schmidtea mediterranea*, *Dugesia japonica* and Mollusca *Lottia gigantean*. Phylogenetic analysis of DBD sequences showed that all of these 2DBD-NRs belonged to a monophyletic group and suggested that 2DBD-NRs originated from a common ancestor gene [8]. Recently, 2DBD-NRs were only identified and/or isolated in Platyhelminths [6, 8–15] and in Mollusca [16, 17]. Until now, there are very few studies on 2DBD-NRs. Our study showed that Sm2DBD-NRα could form a homodimer but could not form a heterodimer with RXRs [8]. By searching structurally homologous sequences in the protein data bank (PDB), Alvite et al. showed that unsaturated fatty acids are preferred ligands by a *Echinococcus granulosus* 2DBD-NR (Eg2DBDa.1) [9]. Tarp et al. showed that *S. mediterranea* 2DBD-NR (nhr-1) was only detected in male and female accessory reproductive organs, and they suggested that *S. mediterranea* 2DBD-NR (nhr-1) was required for planarian reproductive maturation [12].

In this study, 2DBD-NRs are mined from different databases and are phylogenetically analyzed. From our phylogenetic analysis of 2DBD-NRs and following the rule of Nomenclature System for the Nuclear Receptors, a nomenclature for 2DBD-NRs is proposed.

## MATERIALS AND METHODS

### 1. Data mining

2DBD-NRs were mined from The National Center for Biotechnology Information (NCBI) protein database (https://www.ncbi.nlm.nih.gov/), the Ensembl Genomes project *Capitella_teleta* database (https://metazoa.ensembl.org/Capitella_teleta/Tools/Blast) [18], *Notospermus geniculatus* database (https://marinegenomics.oist.jp/nge_v2/blast/search?project_id=52) and *Phoronis australis database* (https://marinegenomics.oist.jp/pau_v2/viewer/info?project_id=51) [19]. Amino acid sequences of both DBDs of Sm2DBD-NRα (AH013462) and Cg2DBD-NR (XP_019919868) were used as the query to pblast (with E-value threshold: 1e-1) against all available NCBI protein databases, and tblastn (with E-value threshold: 1e-1) against *C. teleta*, *N. geniculatus* and *P. australis* genome databases. Any sequence that contains a zinc finger structure of the DBD of NRs (Cys-X2-Cys-X13-Cys-X2-Cys or Cys-X5-Cys-X9-Cys-X2-Cys) was retained (Fig. 1B). After careful check by eye, all amino acid sequences containing two DBDs, partial two DBDs or highly conserved sequence to 2DBD-NRs were retained.

### 2. Phylogenetic analysis

Phylogenetic trees of 2DBD-NRs were constructed from deduced amino acid sequences of both the first and the second DBDs. The amino acid sequences are aligned with ClustalW [20], phylogenetic analysis of the data set is carried out using Bayesian inference MrBAYES v3.1.1 [21] as in our previous study [15]. Only Bayesian inference was carried out in this study because our previous study demonstrated that Bayesian inference highly supports phylogenetic analysis of NRs more than other methods [15]. The trees were started randomly with a mix amino acid replacement model + gamma rates. Two sets of four simultaneous Markov chains were run for 5 million generations and the trees were sampled every100 generations. The Bayesian posterior probabilities (BPPs) were calculated using a Markovchain Monte Carlo (MCMC) sampling approach implemented in MrBAYES v3.1.1 sequences of and the burn-in value was set at 12,500.

### 3. Amino acid sequence analysis

Amino acid sequences of every 2DBD-NRs including full length or partial sequences were aligned using ClustalW [20] and the conserved sequences were identified. Sequence Logos were created online (https://weblogo.berkeley.edu/logo.cgi) [22].

## RESULTS AND DISSCUSSION

### 1. Identification and phylogenetic analysis of 2DBD-NRs

2DBD-NRs were identified in different animal species including those from protostome Spiralia (Rotifera, Brachiopoda, Mollusca, Annelida, Platyhelminthes, Brachiopoda, Nemertea and Phoronida) and Ecdysozoa (Nematoda), and from deuterostome Unchordata (Echinodermata and Chordata (Cephalochordata). Previously, 2DBD-NR were thought to be lophotrochozoan-specific [12], this study shows that 2DBD-NRs broadly exist in protostome and deuterostome species, this further indicates that 2DBD-NR gene was already present in the urbilaterian, a common ancient ancestor of protostomes and deuterostomes.

Phylogenetic analysis of identified 2DBD-NRs using amino acid sequence of both the first and the second DBD was carried out by Bayesian inference. The result shows that all of the 2DBD-NRs from protostome Spiralia and deuterostomes are clustered together forming a Spiralia/deuterostomes group (Bayesian posterior probabilities (BPPs) = 0.98), while all Ecdysozoa Nematoda 2DBD-NRs are clustered outside of the Spiralia/deuterostomes group. This result suggests that protostome Spiralia and deuterostomes share a close common ancestor 2DBD-NR gene (Fig. 2). In protostome spiralia/deuterostomes group, 2DBD-NRs are clustered in two groups: 2DBD-NRA and 2DBD-NRB with BBP = 0.97 and 1, respectively (Fig. 2). Both 2DBD-NRA and 2DBD-NRB groups contain members from protostome and deuterostome species, this result suggests that two ancient 2DBD-NR genes (*2DBD-NRA* and *2DBD-NRB*) were present in a common ancestor of protostomes and deuterostomes. Phylogenetic analysis further shows that most 2DBD-NRs gene underwent duplications after the split of different animal phyla. Thus, 2DBD-NR in a certain animal phylum may be a paralogue, rather than an orthologue, of that in another animal phylum (Fig. 2). For example, Rotifera 2DBD-NRs are clustered in five subgroups in the 2DBD-NRA group, and another four subgroups are clustered together without members from any other Phylum.

**Figure 2.**
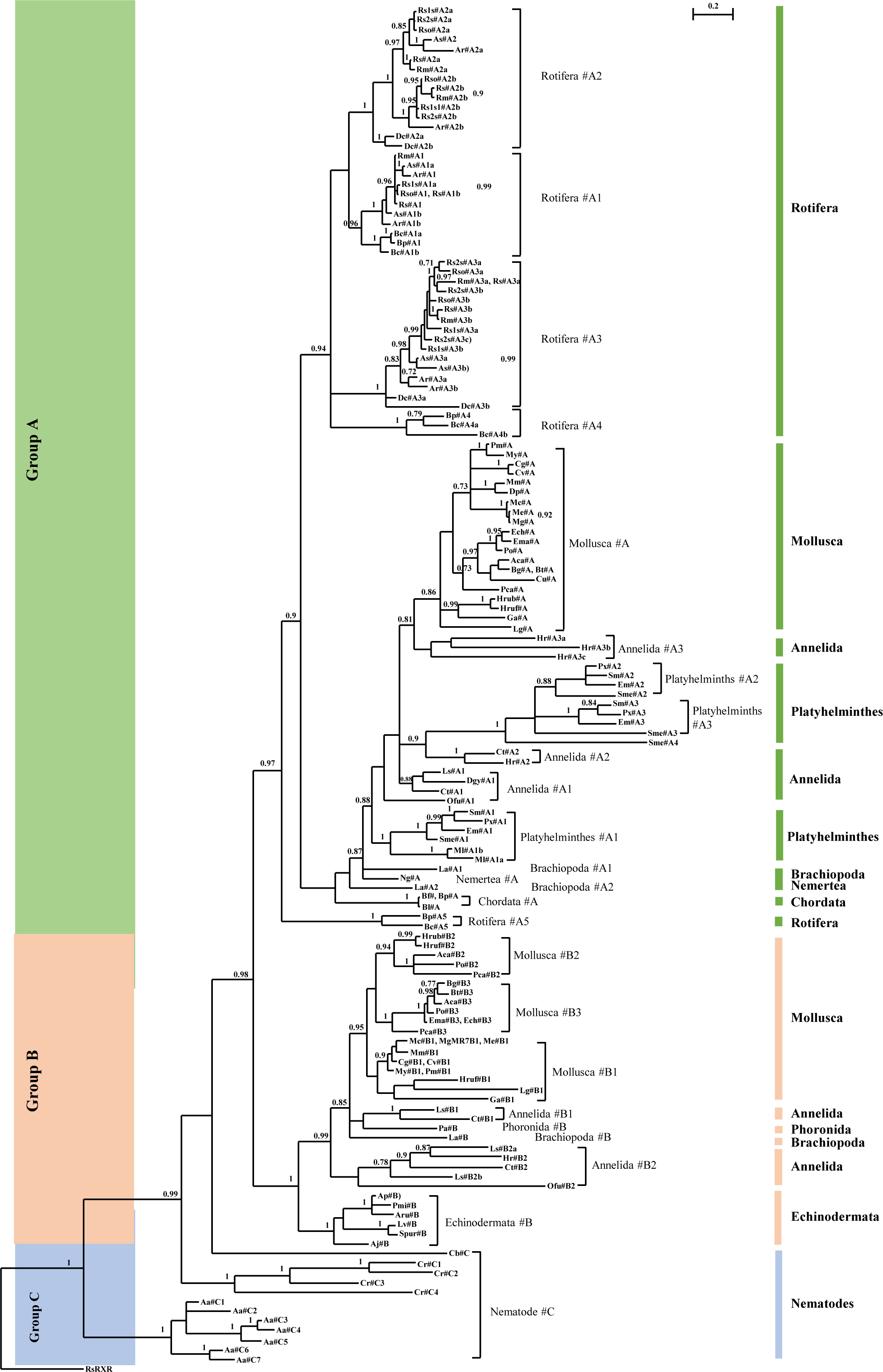
Bayesian phylogenetic analysis of 2DBD-NRs. Bayesian phylogenetic tree is constructed with the amino acid sequence of both the first and second DBD. The BPPs values are shown above each branch or after the name of the NR, branches under the PPs 0.5 are shown as polytomies. **Aa**: *Aphelenchus avenae*, **Aca**: *Aplysia californica*, **Aj**: *Anneissia japonica*, **Ap**: *Acanthaster planci*, **Ar**: *Adineta ricciae*, **Aru**: *Asterias rubens*, **As**: *Adineta steineri*, **Bc**: *Brachionus calyciflorus*, **Bf**: *Branchiostoma floridae*, **Bg**: *Biomphalaria glabrata*, **Bl**: *Branchiostoma lanceolatum*, **Bt**: *Bulinus truncatus*, **Bp**: *Brachionus plicatilis*, **Cb**: *Caenorhabditis brenneri*, **Cg**: *Crassostrea gigas*, **Cr**: *Caenorhabditis remanei*, **Cu**: *Candidula unifasciata*, **Cv**: *Crassostrea virginica*, **Ct**: *Capitella teleta*, **Dc**: *Didymodactylos carnosus*, **Dgy**: *Dimorphilus gyrociliatus*, **Dp**: *Dreissena polymorpha*, **Ech**: *Elysia chlorotica*, **Em**: *Echinococcus multilocularis*, **Ema**: *Elysia marginata*, **Ga**: *Gigantopelta aegis*. **Hr**: *Helobdella robusta*, **Hrub**: *Haliotis rubra*, **Hruf**: *Haliotis rufescens*, **La**: *Lingula anatina*, **Lg**: *Lottia gigantea*, **Ls**: *Lamellibrachia satsuma*, **Lv**: *Lytechinus variegatus*, **Mco**: *Mytilus coruscus*, **Me**: *Mytilus edulis*, **Mg**: *Mytilus galloprovincialis*, **Ml**: *Macrostomum lignano*, **Mm**: *Mercenaria mercenaria*, **My**: *Mizuhopecten yessoensis*, **Ofu**: *Owenia fusiformis*, **Pa**: *Phoronis australis*, **Pca**: *Pomacea canaliculata*, **Pm**: *Pecten maximus*, **Pmi**: *Patiria miniata*, **Po**: *Plakobranchus ocellatus*, **Px**: *Protopolystoma xenopodis*, **Rm**: *Rotaria magnacalcarata*, **Rs**: *Rotaria socialis*, **Rso**: *Rotaria sordida*, **Rs1s**: *Rotaria sp. Silwood1*, **Rs2s**: *Rotaria sp. Silwood2*, **Sm**: *Schistosoma mansoni*, **Sme**: *Schmidtea mediterranea*, **Spur**: *Strongylocentrotus purpuratus*. **#**: 2DBD-NR. The capital letter (A, B or C) after # (2DBD-NR) indicates 2DBD-NR group, Arabic numeral after capital letter indicates individual gene, and a lowercase letter at the end of the gene indicates variant. *: All member of Rotifers 2DBD-NRA3 and 2DBD-NRA5 groups, **: All member of Echinodermata 2DBD-NRB group. GenBank Accession number of analyzed NRs see Table 6.

### 2. 2DBD-NRs in different animals

Members from both 2DBD-NRA and 2DBD-NRB groups are identified in Mollusca, Annelida and Brachiopoda; while 2DBD-NRA is not found in Phoronida and Echinodermata, 2DBD-NRB is not identified in Platyhelminthes, Nemertea, Rotifera and Chordata. Since most 2DBD-NRs are paralogues in different animal phylum, we present our findings below by animal phyla.

**1) 2DBD-NRs in Mollusca.** 2DBD-NRs are identified in Mollusca species from Class Bivalvia and Class Gastropoda (Table 1). Phylogenetic analysis shows that Mollusca 2DBD-NRs are clustered in both 2DBD-NRA and 2DBD-NRB groups. 2DBD-NRA group contains only one member from each analyzed Mollusca species, it suggests that one 2DBD-NRA is present in Mollusca species from Class Bivalvia and Class Gastropoda. Mollusca 2DBD-NRB group contains three subgroups (2DBD-NRB1, 2DBD-NRB2 and 2DBD-NRB3), it suggests that three 2DBD-NRBs are present in analyzed Mollusca species (Fig. 2). All of the three 2DBD-NRBs (2DBD-NRB1, 2DBD-NRB2 and 2DBD-NRB3) are found in species from Gastropoda, but only one 2DBD-NRB (2DBD-NRB1) is identified in species from Class Bivalvia. This result suggests that four 2DBD-NRs (2DBD-NRA, 2DBD-NRB1, 2DBD-NRB2 and2DBD-NRB3) are present in Gastropoda and two 2DBD-NRs (2DBD-NRA and 2DBD-NRB1) are present in Bivalvia. Since Mollusca 2DBD-NRB2 and 2DBD-NRB3 groups share a shallower node, it suggests that 2DBD-NRB2 and 2DBD-NRB3 were formed by recent gene duplication (Fig. 2 and Table 1).

**Table 1.**
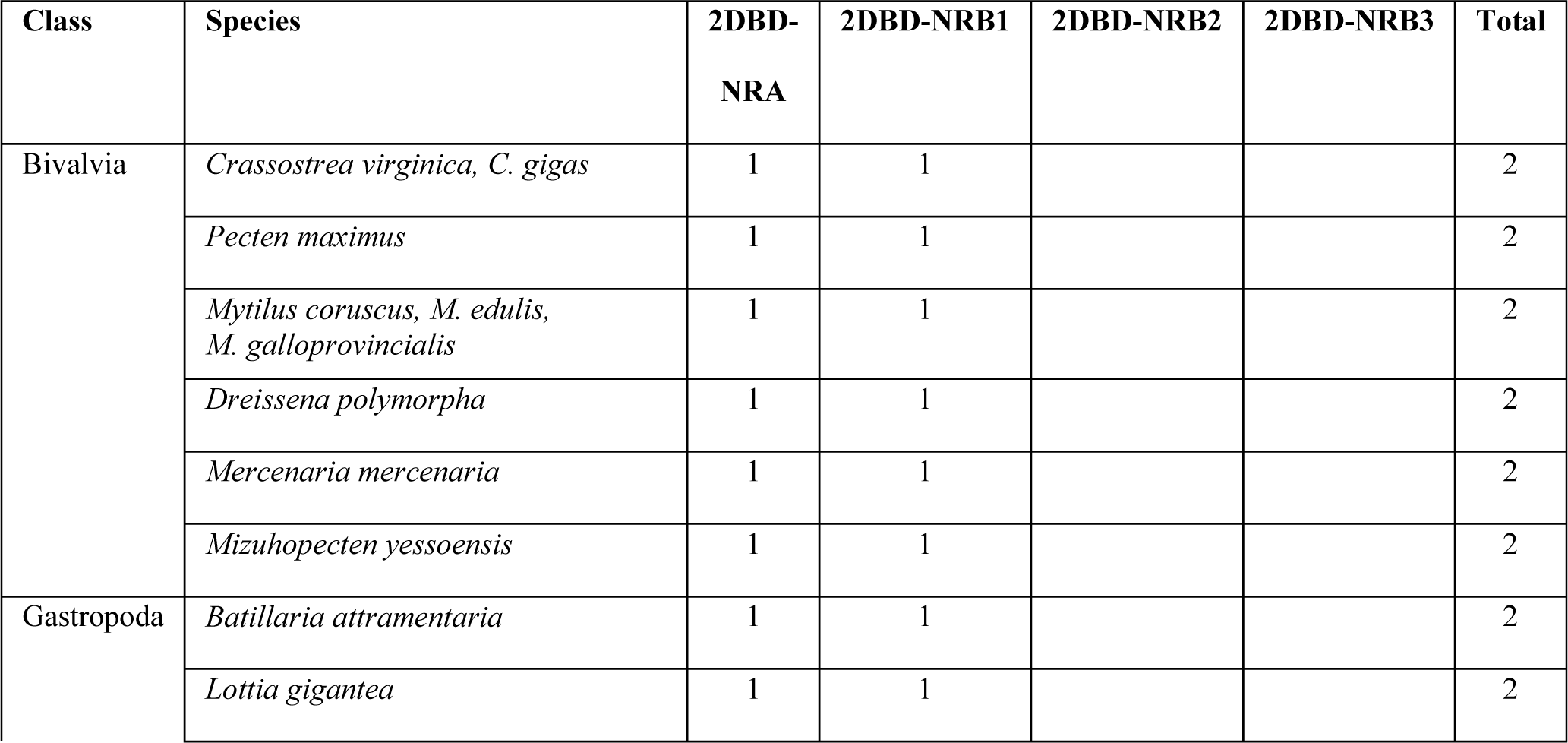
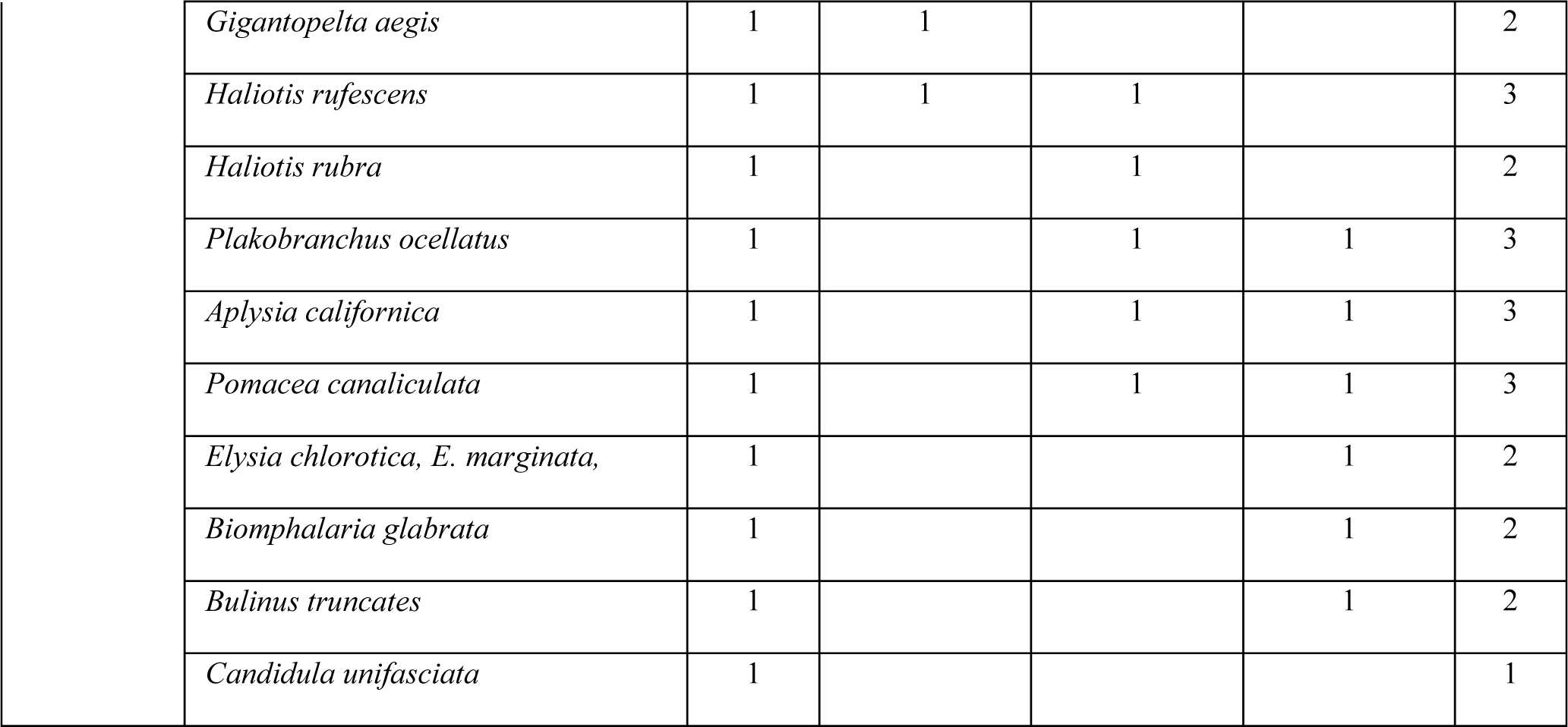
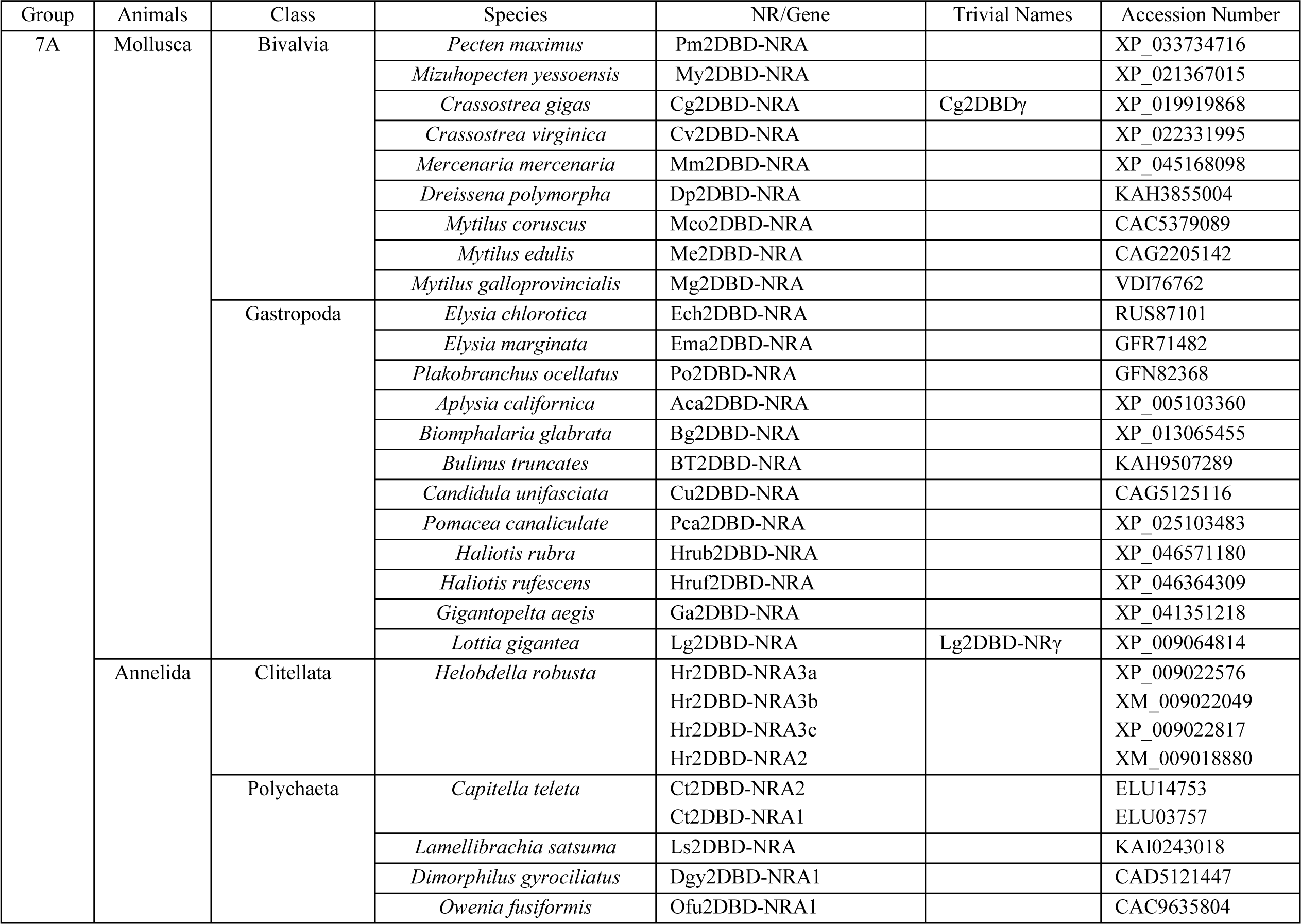
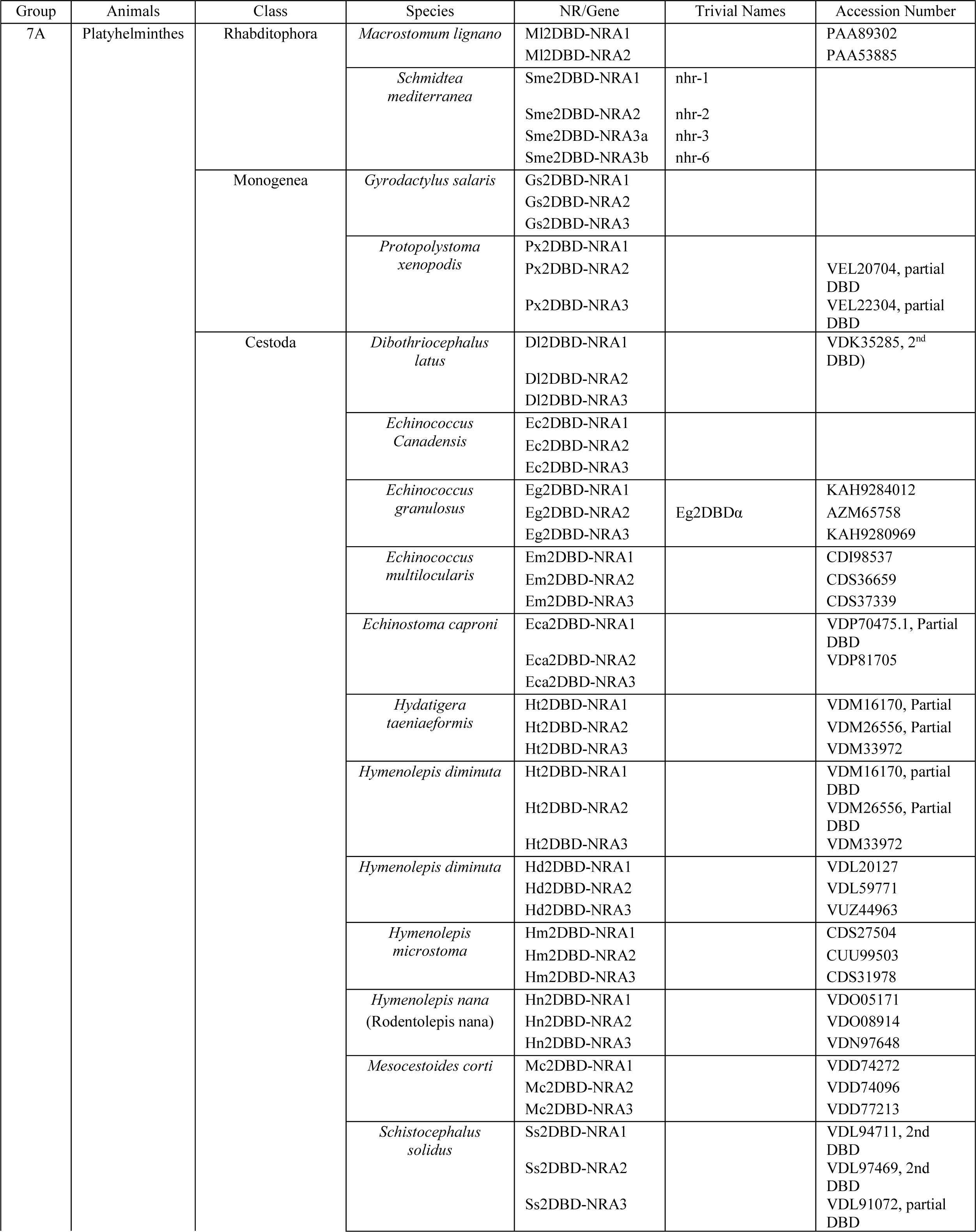
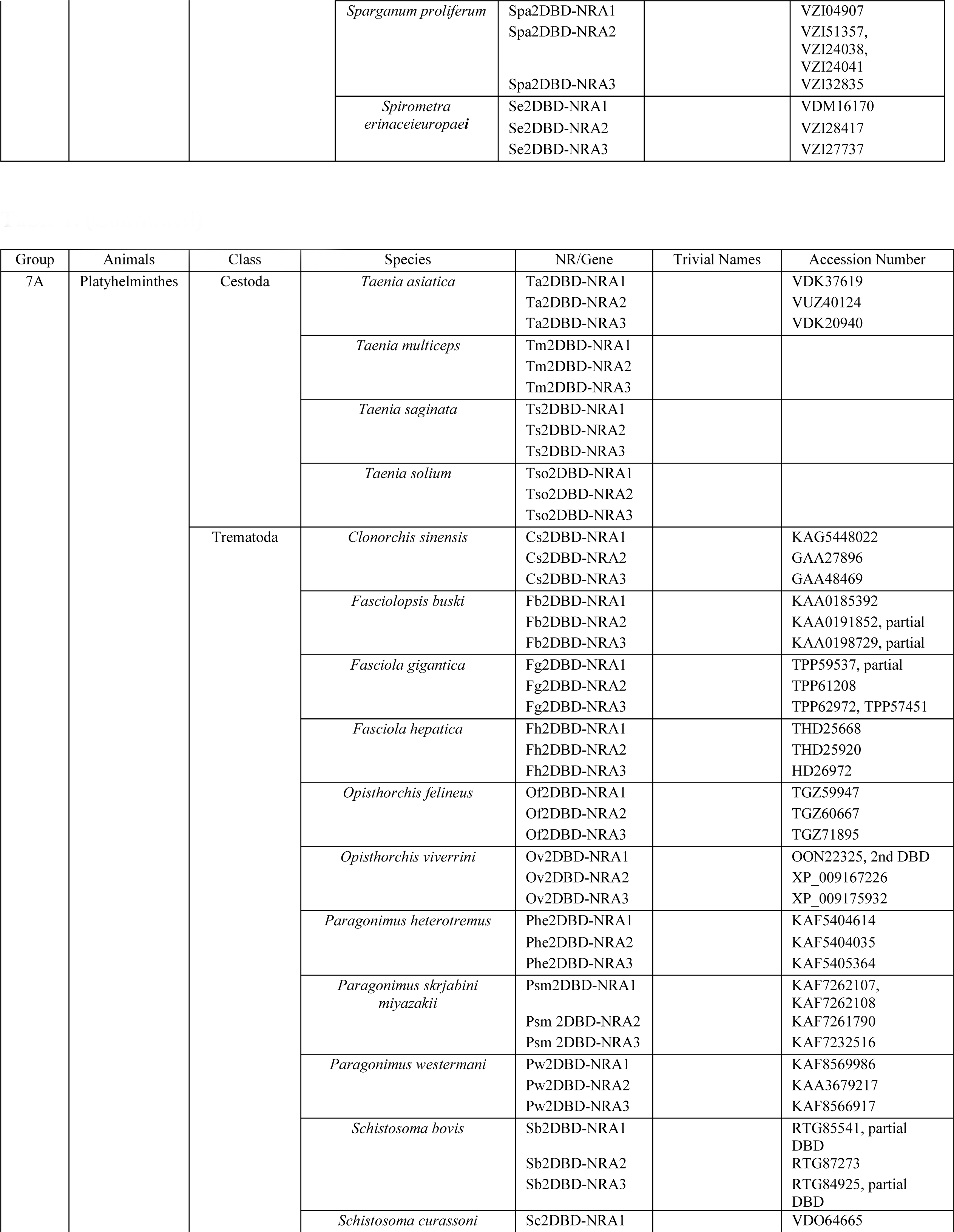
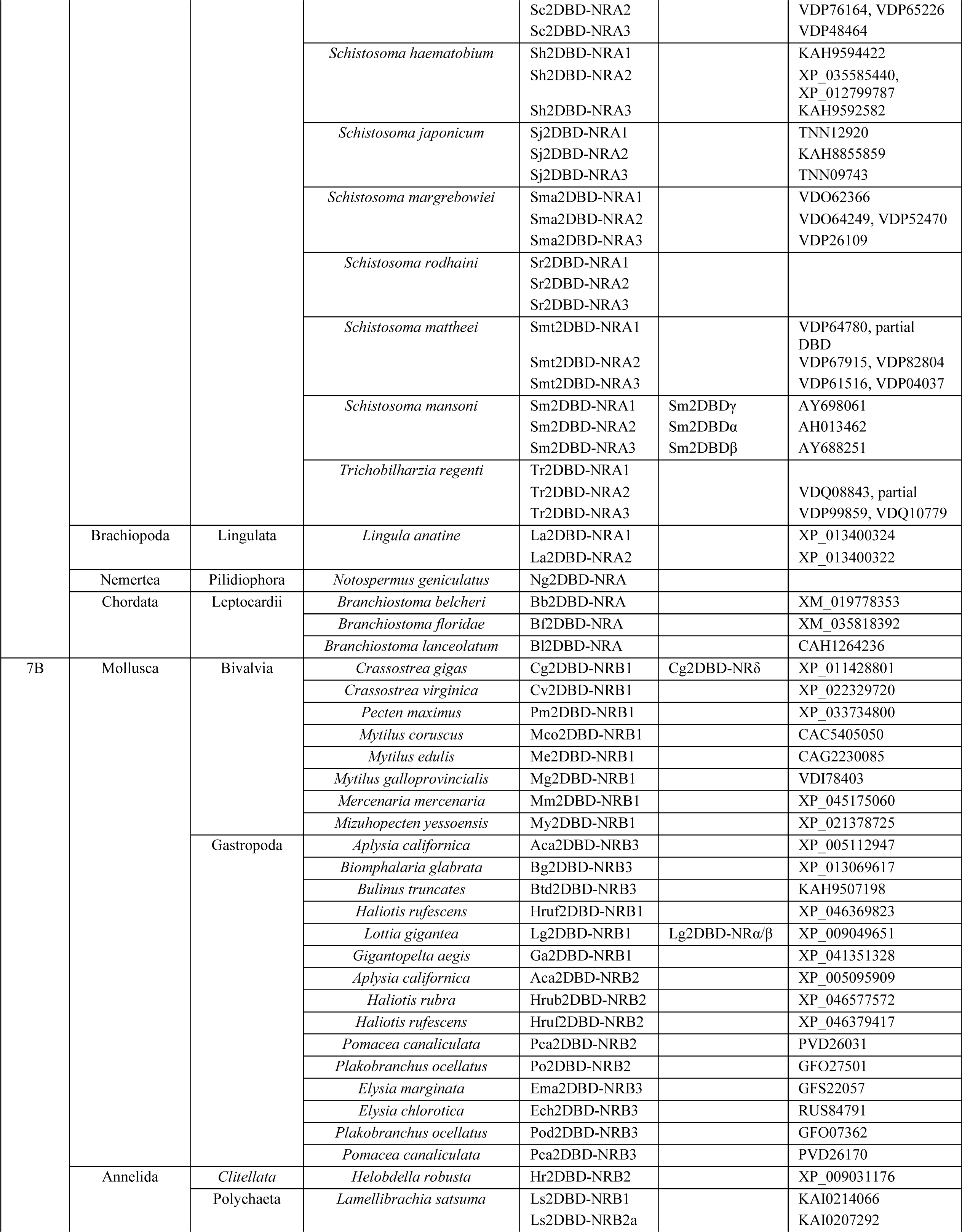
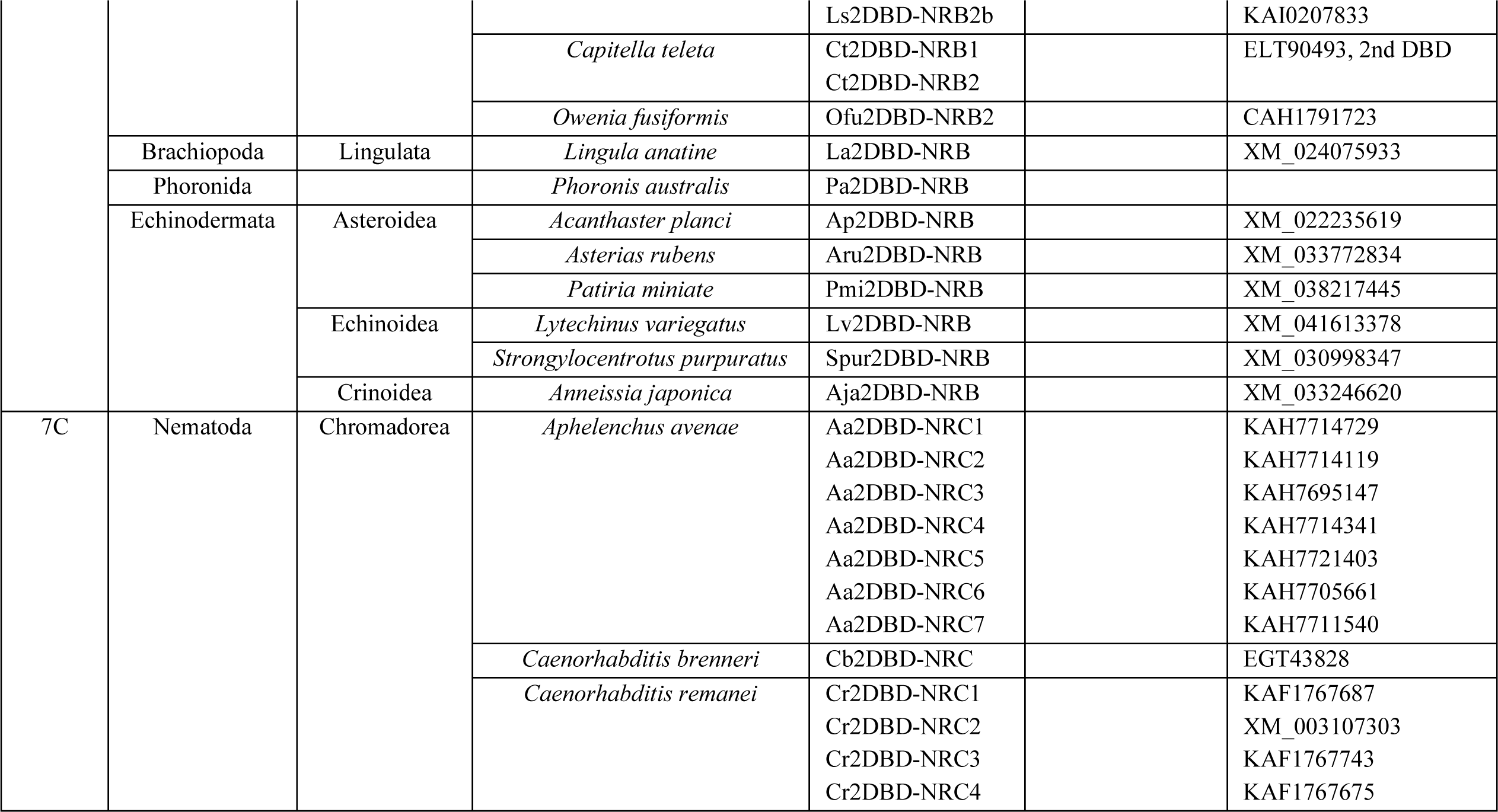
2DBD-NRs identified in Mollusca.

Mining genome database of Mollusca *Crassostrea gigas*, Vogeler et al. [17] identified two *C. gigas* 2DBD-NRs (Cg2DBDγ and Cg2DBDδ). They showed that Cg2DBDγ contained the same unique P-box sequences, CEACKK, as *S. mansoni* 2DBD-NRs in the first DBD, but Cg2DBDδ contained a different P-box sequence, CLPCKS, in the first DBD, this P-box sequence was not found in any other nuclear receptor. Amino acid sequence alignment shows that 2DBD-NRBs from Annelida, Brachiopoda and Phoronida and Mollusca possess the same P-box sequence CLPCKS in their first DBD with few members demonstrating divergent sequences (Supplemental material 1).

**2) 2DBD-NRs in Brachiopoda.** Three members are identified in Brachiopoda *Lingula anatine*.

MrBayes inference shows that two of them belong to 2DBD-NRA group (La2DBD-NRA1 and La2DBD-NRA2) and one is clustered in 2DBD-NRB group (La2DBD-NRB) (Fig. 2).

**3) 2DBD-NRs in Phoronida.** One member is identified in Phoronida *Phoronis australis* that belongs to 2DBD-NRB group (Pa2DBD-NRB) (Fig. 2).

**4) 2DBD-NRs in Nemertea.** One member is identified in Nemertea *Notospermus geniculatus* and it is clustered in 2DBD-NRA group (Ng2DBD-NRA) (Fig. 2).

**5) 2DBD-NRs in Annelida.** 2BDB-NRs are identified in Annelida species from Class Polychaeta and Class Clitellata, they are clustered in five subgroups. Three subgroups are clustered in 2DBD-NRA group and two subgroups are in the 2DBD-NRB group. This result suggests that at least five 2DBD-NRs exist in Annelida. 2DBD-NRA2 and 2DBD-NRB2 are identified in both Class Polychaeta and Clitellata, 2DBD-NRA1 and 2DBD-NRB1 are identified only in Class Polychaeta, while 2DBD-NRA3 is only identified in Class Clitellata (Fig. 2, Table 2).

**Table 2.**
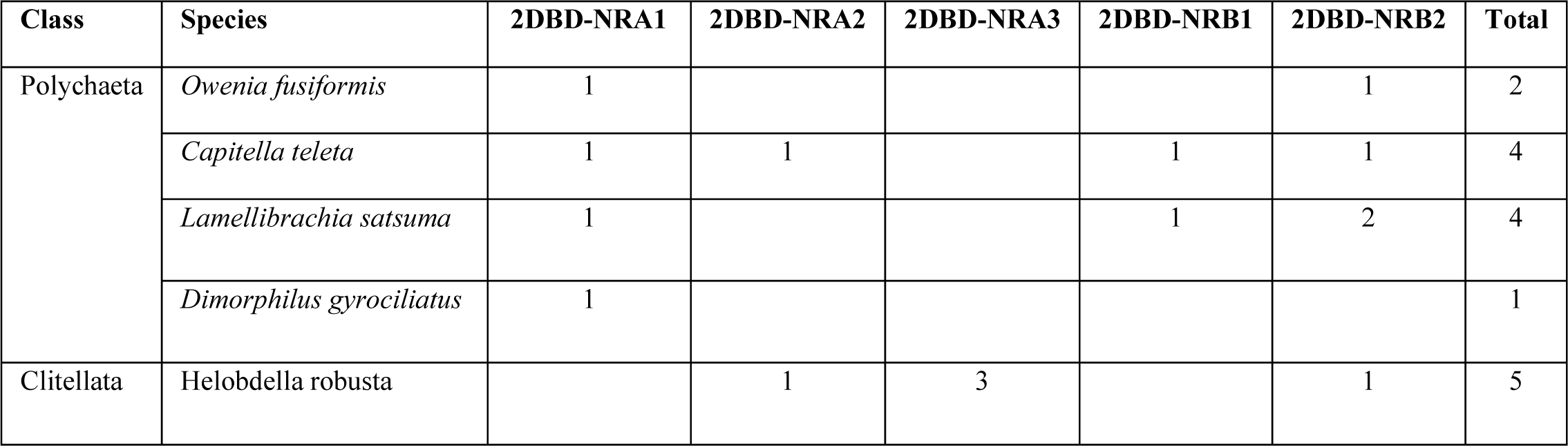
2DBD-NRs in Annelida.

**6) 2DBD-NRs in Rotifera.** NRs were found genome-wide in Rotifera in different species of *Brachionus*, no 2DBD-NR were identified from Rotifera in a previous study [23]. In this study, 2DBD-NRs are identified in ten species from two subclasses (Bdelloidea and Monogononta) of Class Eurotatoria (Table 3) in Rotifera. These 2DBD-NRs are clustered in five subgroups in 2DBD-NRA group. This result suggests that five 2DBD-NRAs are present in Rotifera Eurotatoria Class. 2DBD-NRA1 are identified in both subclasses, 2DBD-NRA2 and 2DBD-NRA3 are only identified in subclass Bdelloidea, while 2DBD-NRA4 and 2DBD-NRA5 are only identified in subclass Monogononta. The significant character of Rotifera 2DBD-NRs is that two or more divergent copies of every 2DBD-NR gene are present, this is consistence with the previous study of the genome of Bdelloidea [24, 25] (Fig. 2 and Table 3).

**Table 3.** 2DBD-NRs in Rotifera.

| Class | Subclass | Species | 2DBD-NRA1 | 2DBD-NRA2 | 2DBD-NRA3 | 2DBD-NRA4 | 2DBD-NRA5 | Total |
| --- | --- | --- | --- | --- | --- | --- | --- | --- |
| Eurotatoria | Bdelloidea | *Rotaria magnacalcarata* | 1 | 2 | 2 |  |  | 5 |
|  |  | *Adineta steineri* | 2 | 1 | 2 |  |  | 5 |
|  |  | *Adineta ricciae* | 2 | 2 | 2 |  |  | 6 |
|  |  | *Rotaria sp. Silwood1* | 1 | 3 | 2 |  |  | 6 |
|  |  | *Rotaria sp. Silwood2* |  | 2 | 3 |  |  | 5 |
|  |  | *Rotaria sordida* | 1 | 2 | 2 |  |  | 5 |
|  |  | *Rotaria socialis* | 1 | 2 | 2 |  |  | 5 |
|  |  | *Didymodactylos carnosus* |  | 2 | 2 |  |  | 4 |
|  | Monogononta | *Brachionus calyciflorus* | 2 |  |  | 2 | 1 | 5 |
|  |  | *Brachionus plicatilis* | 1 |  |  | 1 | 1 | 3 |

**7) 2DBD-NRs in Platyhelminths.** Our previous study showed that two 2DBD-NRs were present in Rhabditophora, *Macrostomum lignano* [10, 15] and four in *Schmidtea mediterranea*. Three members were present in parasitic species from Platyhelminths including Class Monogenea, Cestoda and Trematoda, respectively [6, 8, 9, 11-15]. Phylogenetic analysis of Platyhelminths 2DBD-NRs in this study shows that all Platyhelminths 2DBD-NRs are clustered in 2DBD-NRA group, which suggests that 2DBD-NRB gene is missing in Platyhelminths. All parasitic 2DBD-NRs are clustered in three subgroups: 2DBD-NRA1 (*Schistosoma mansoni* 2DBD-NRγ orthologues), 2DBD-NRA2 (*S. mansoni* 2DBD-NRα orthologues) and 2DBD-NRA3 (*S. mansoni* 2DBD-NRβ orthologues), each subgroup contains members from species of each Class of Monogenea, Cestoda and Trematoda. For free-living flatworms, both *M. lignano* 2DBD-NRs are clustered in Platyhelminths 2DBD-NRA1 subgroup. For the four *S. mediterranea* 2DBD-NRs, one is clustered in Platyhelminths 2DBD-NRA1 subgroup, one is in Platyhelminths 2DBD-NRA2 subgroup and one is clustered in Platyhelminths 2DBD-NRA3 subgroup. The fourth *S. mediterranea* 2DBD-NR (Sme2DBD-NRA4) is on the base of 2DBD-NRA2 and 2DBD-NRA3 subgroups (Fig. 2). This result suggests that 2DBD-NR underwent another round of duplication in a common ancestor of *S. mediterranea* and parasitic Platyhelminths and then one 2DBD-NR was lost in a common ancestor of parasitic Platyhelminths. This result is consistent with our previous study [15]. In this study, 2DBD-NRs are identified in more parasitic Platyhelmins including the lung fluke *Paragonimus westermani, P. skrjabini miyazakii* and *P. heterotremus*, liver fluke *Fasciola gigantica*, giant intestinal flukes *Fasciolopsis buski* and tapeworms *Sparganum proliferum*. Sequence alignment shows that all of them are highly conserved with known Platyhelminths 2DBD-NRs.

**8) 2DBD-NRs in Echinodermata.** One member is identified in each analyzed species of Echinodermata including those from Class Asteroidea, Class Echinoidea and Class Crinoidea. MrBayes inference analysis shows that all Echinodermata 2DBD-NRs form a monophyletic subgroup in 2DBD-NRB group (Fig. 2 and Table 4). Previously, 33 NRs were identified in the genome database of *Strongylocentrotus purpuratus*, but until this report no 2DBD-NR was reported [26].

**Table 4.** 2DBD-NRs in Deuterostome.

| Phyla | Class | Species | 2DBD-NRA | 2DBD-NRB |
| --- | --- | --- | --- | --- |
| Echinodermata | Asteroidea | *Acanthaster planci* |  | 1 |
|  |  | *Asterias rubens* |  | 1 |
|  |  | *Patiria miniate* |  | 1 |
|  | Echinoidea | *Lytechinus variegatus* |  | 1 |
|  |  | *Strongylocentrotus purpuratus* |  | 1 |
|  | Crinoidea | *Anneissia japonica* |  | 1 |
| Chordata | Leptocardii | *Branchiostoma belcheri* | 1 |  |
|  |  | *Branchiostoma floridae* | 1 |  |
|  |  | *Branchiostoma lanceolatum* | 1 |  |

**9) 2DBD-NRs in Chordata.** One member is identified in each species of analyzed Chordata Amphioxi including *Branchiostoma belcheri*, *B. floridae* and *B. lanceolatum*. MrBayes inference analysis shows that all of them form a monophyletic subgroup in 2DBD-NRA group (Fig. 2 and Table 4). Though 33 NRs were identified in *B. floridae,* but until this report no 2DBD-NR was reported in Amphioxi [27],

**10) 2DBD-NRs in Nematoda.** 2DBD-NRs are present in nematode species *Aphelenchus avenae*, *Caenorhabditis brenneri* and *Caenorhabditis remanei*. Four 2DBD-NRs are found in *C. remanei*, one in *C. brenneri* and seven in *A. avenae*. Phylogenetic analysis shows that all these 2DBD-NRs are clustered outside of Spiralia/deuterostomes 2DBD-NR group (Fig. 2). Sequence alignment of DBDs shows that the P-box sequence is highly divergent in Nematoda 2DBD-NRs (Supplemental material 1). These results suggests that Nematode 2DBD-NRs underwent extensive divergence as known in other nematode NRs [28–31].

See Table 5 for a summary of the numbers of 2DBD-NRs identified in different animal species.

**Table 5.**
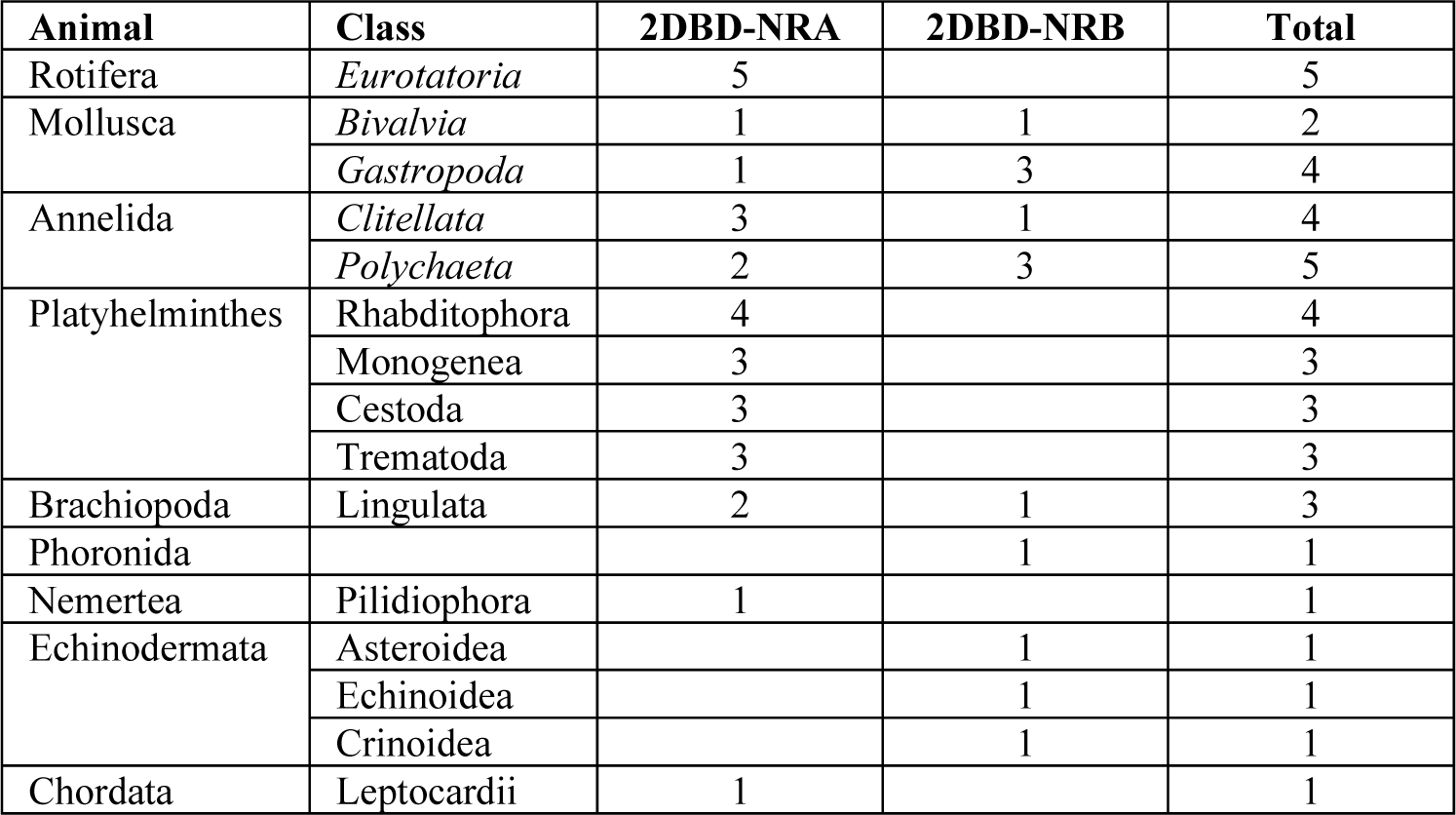
Numbers of 2DBD-NRs identified in different animals.

### 3. Amino acid sequence analysis of 2DBD-NRs

**1) AB domain.** A highly conserved ‘N-terminal signature sequence’ (NTSS [32]) is found at the 3’end of the A/B domain of parasitic Platyhelminths 2DBD-NRs. A NTSS of CNLGXKDRRP is present in Platyhelminth Trematode 2DBD-NRA1s, and a NTSS of TNDVTAMKEKTP is present in Cestoda 2DBD-NRA1s. A NTSS of (S/T)PEXAFXQYQXR(M/S)EGQX represents both Platyhelminths 2DBD-NRA2s and 2DBD-NRA3s (Fig. 3). That the fact that Platyhelminth 2DBD-NRA2 and 2DBD-NRA3 members share a conserved NTSS further supports our phylogenetic analysis.

**Figure 3.**
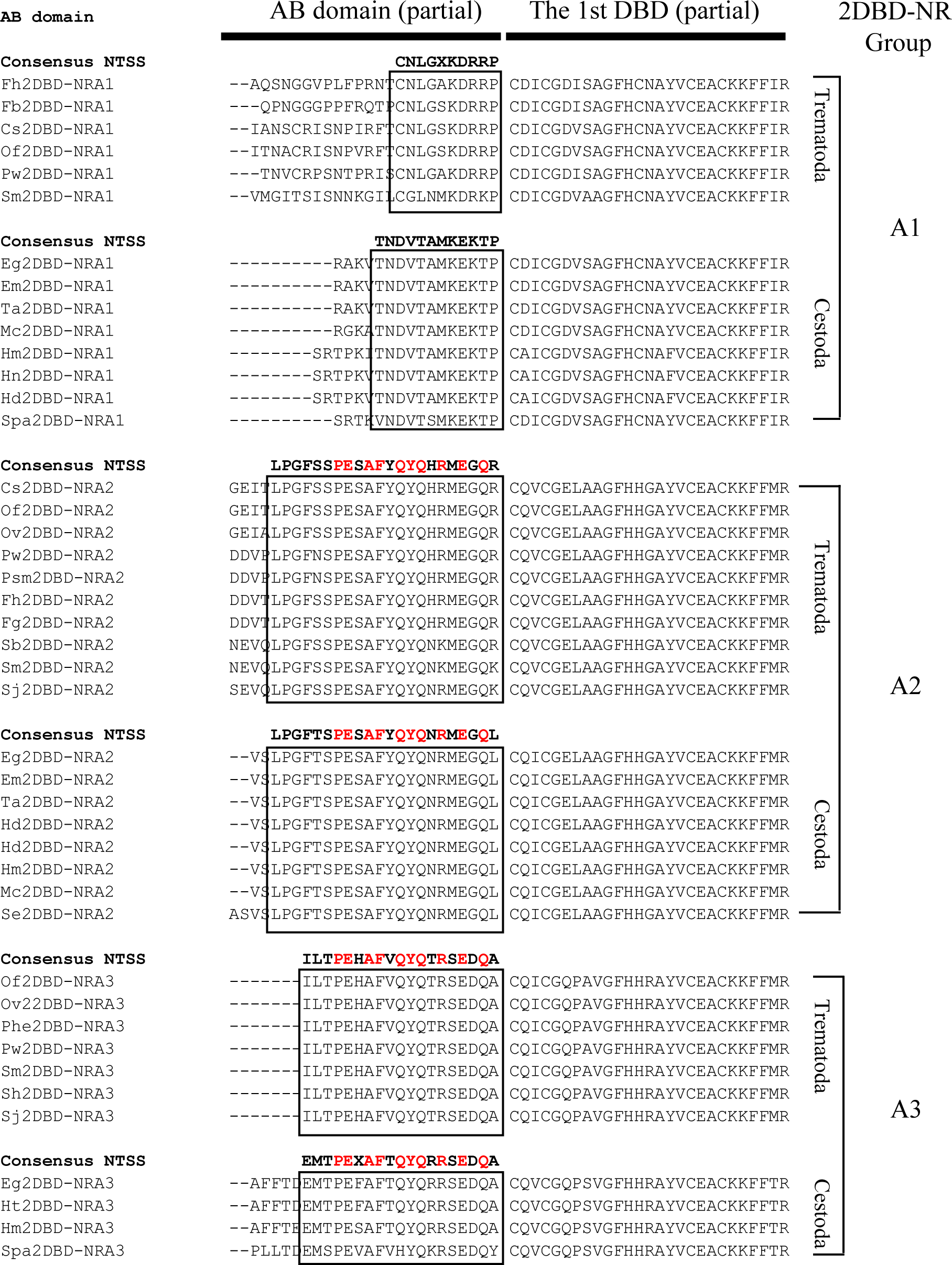
Amino acid sequence alignment shows the conserved NTSS in parasitic Platyhelminths 2DBD-NRs. **Cs**: *Clonorchis sinensis*, **Eg**: *Echinococcus granulosus*, **Em**: *Echinococcus multilocularis*, **Fb**: *Fasciolopsis buski*, **Fh**: *Fasciola hepatica,* **Fg**: *Fasciola gigantica*, **Hd**: *Hymenolepis diminuta*, **Hm**: *Hymenolepis microstoma*, **Hn**: *Hymenolepis nana*, **Ht**: *Hydatigera taeniaeformis*, **Of**: *Opisthorchis felineus*, **Ov**: *Opisthorchis viverrini*, **Phe**: *Paragonimus heterotremus*, **Mc**: *Mesocestoides corti*, **Pw**: *Paragonimus westermani*, **Psm**: *Paragonimus skrjabini miyazakii*, **Se**: *Spirometra erinaceieuropaei*, **Sb**: *Schistosoma bovis*, **Sh**: *Schistosoma haematobium*, **Sj**: *Schistosoma japonicum*, **Spa**: *Sparganum proliferum*, **Sm**: *Schistosoma mansoni*, **Ta**: *Taenia asiatica*. The red letters indicate the amino acid residues that are conserved in parasitic Platyhelminths 2DBD-NRA2 and 2DBD-NRA3, it suggests that parasitic Platyhelminths 2DBD-NRA2 and 2DBD-NRA3 underwent recent duplication.

**2) DBDs.** DBD is the most conserved region in NRs, this region contains two highly conserved C4 type zinc fingers with a module of C-X2-C-X13-C-X2-C for the first Zinc finger (CI) and C-X5-C-X9-C-X2-C for the second Zinc finger (CII), where C represents cysteine, X represents any variable and the next number indicates the number of amino acids between the cysteines [33]. Amino acid alignment of all identified 2DBD-NRs in this study shows that both DBDs of most of the 2DBD-NRs contain the conserved zinc finger modules as above with very little exception (Fig. 1B and Supplemental material 2). Recently, we reported a novel zinc finger CHC2 motif (C-X6-C-X9-H-X2-C) in DBD of parasitic Platyhelminth NRs [15], this motif is not identified in any 2DBD-NRs.

In 1989, Umesono and Evans [34] identified two conserved motifs in DBD of NRs. One motif was defined as proximal box (P-box), it was followed the third Cysteine of zinc finger I (CI) including five amino acids. The other motif was distal box (D-box) which was located between the fifth and sixth Cysteine in zinc finger II (CII) of DBD (Fig. 1B). The P-box is critical for identifying the primary nucleotide sequence of the half-sites and the D-box is important for protein dimerization. Previously, we demonstrated that 2DBD-NRs possess a unique P-box sequence (EACKK) in the first DBD that is not present in any other NRs. The P-box sequence of the second DBD (EGCKG) followed by the amino acid sequence FFRR (EGCKGFFRR) is identical to that of most members in NR subfamily 1 (NR1) [6, 8]. Recently, Vogeler et al. [17] showed that a unique P-box sequence (LPCKS) was present in the first DBD of a 2DBD-NR in Mollusca *C. gigas* [17]. In this study, various different P-box sequences are found in 2DBD-NRs. The P-box sequence in the first DBD and the second DBD of 2DBD-NRs (**P-P module**) is found to be different among 2DBD-NRAs, 2DBD-NRBs and 2DBD-NRCs (Fig. 1C and Supplemental material 2). The P-P module of 2DBD-NRA is EACKK-EGCKG with a few divergence sequences in the second DBD (Supplemental material 1 and 2). Two types of P-P modules are found in 2DBD-NRB, one is LPCKS-EGCKK, and it is present in 2DBD-NRBs of Mollusca, Annelida, Brachiopoda and Phoronida, with a divergent sequence in the P-box of the first DBD in Annelida and Brachiopoda 2DBD-NRB (Supplemental material 1 and 2). The P-box sequence of LPCKS was first identified in Cg2DBDδ [17]. The P-P module EACKS-EGCKG is only found in Echinodermata 2DBD-NRBs. Interestingly, the P-box sequence of the first DBD of Echinodermata 2DBD-NRBs (EACKS) is identical to that of basal metazoans Porifera NRs (SdRXR and RsNR1) and the P-box sequence of the second DBD is identical to that of most members of 2DBD-NRA, and some members from NR subfamily 1, 2, 4, 6 and 8 (Fig. 1C). The different P-P module among 2DBD-NRs suggests that the mechanism of DNA binding may different in 2DBD-NRAs, 2DBD-NRBs and 2DBD-NRCs. The D-box of NR is located between the fifth and sixth Cysteine in zinc finger II (CII) of DBD with five amino acid residue between the two Cysteines (C-X5-C) [34]. Amino acid sequence alignment of 2DBD-NRs shows that most of 2DBD-NRs have a conserved D box with C-X5-C in both DBDs (Supplemental material 2).

**3) Amino acid sequence between the first and second DBDs.**

Amino acid sequence alignment shows that most member of the 2DBD-NRA and 2DBD-NRB possess 17 - 22 amino acids between two DBDs. Highly conserved sequences in this region are found in parasitic Platyhelminths 2DBD-NRAs (Fig. 4), Rotifera 2DBD-NRAs (Fig. 5) and in 2DBD-NRBs (Fig. 6), respectively.

**Figure 4.**
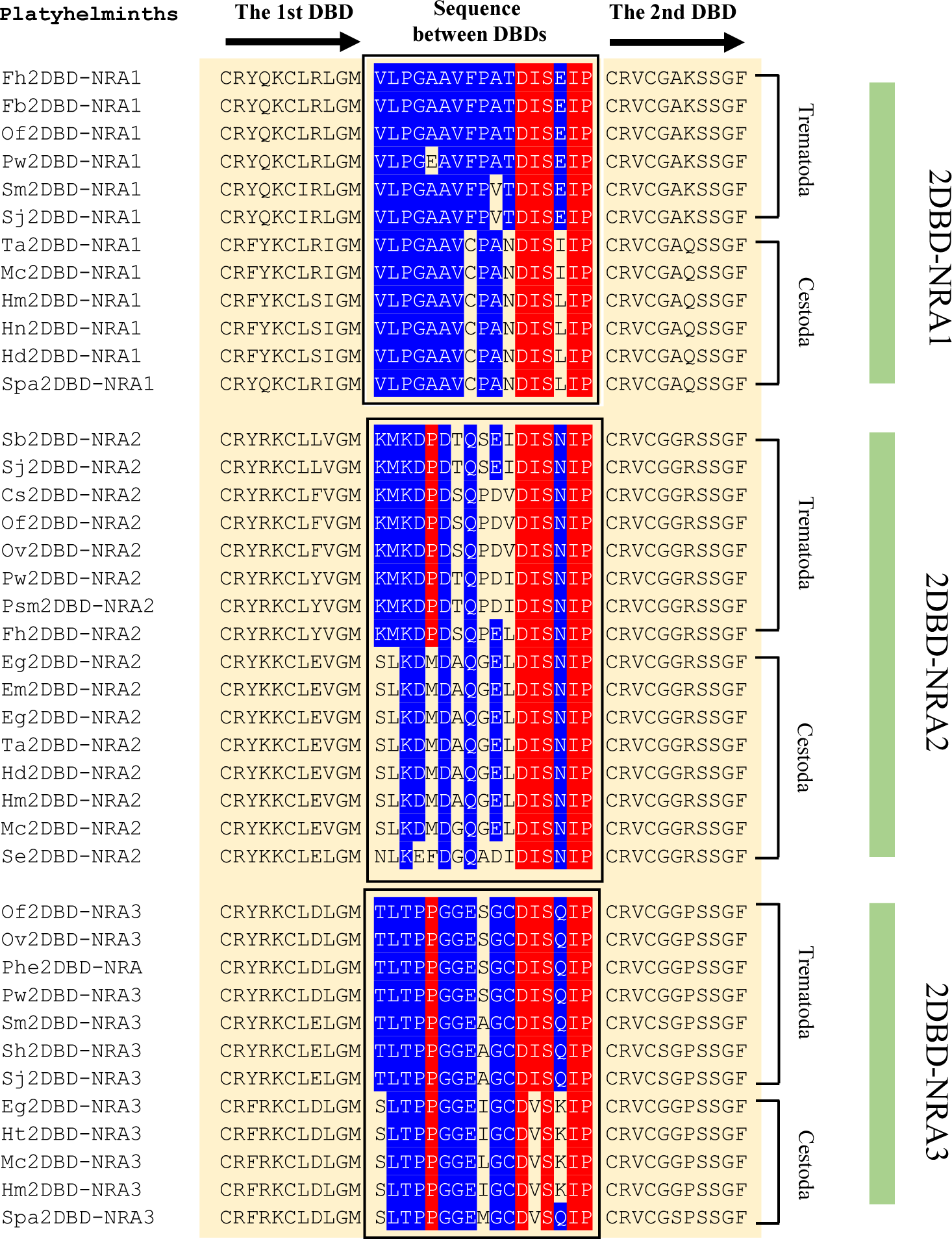
Amino acid sequence between the first and second DBDs in Platyhelminth 2DBD-NRs. Cs: *Clonorchis sinensis*, Eg: *Echinococcus granulosus*, Em: *Echinococcus multilocularis*, Fb: *Fasciolopsis buski*, Fh: *Fasciola hepatica,* Hd: *Hymenolepis diminuta*, Hm: *Hymenolepis microstoma*, Hn: *Hymenolepis nana*, Ht: *Hydatigera taeniaeformis*, Mc: *Mesocestoides corti*, Of: *Opisthorchis felineus,* Ov: *Opisthorchis viverrini*, Pw: *Paragonimus westermani*, Psm: *Paragonimus skrjabini miyazakii*, Sj: *Schistosoma japonicum*, Se: *Spirometra erinaceieuropaei*, Sh: *Schistosoma haematobium*, Sm: *Schistosoma mansoni*, Spa: *Sparganum proliferum*, Ta: *Taenia asiatica*. Blue highlighted letters indicate the conserved amino acid residues in each subgroup and red highlighted letters indicate the conserved amino acid residues among subgroups.

**Figure 5.**
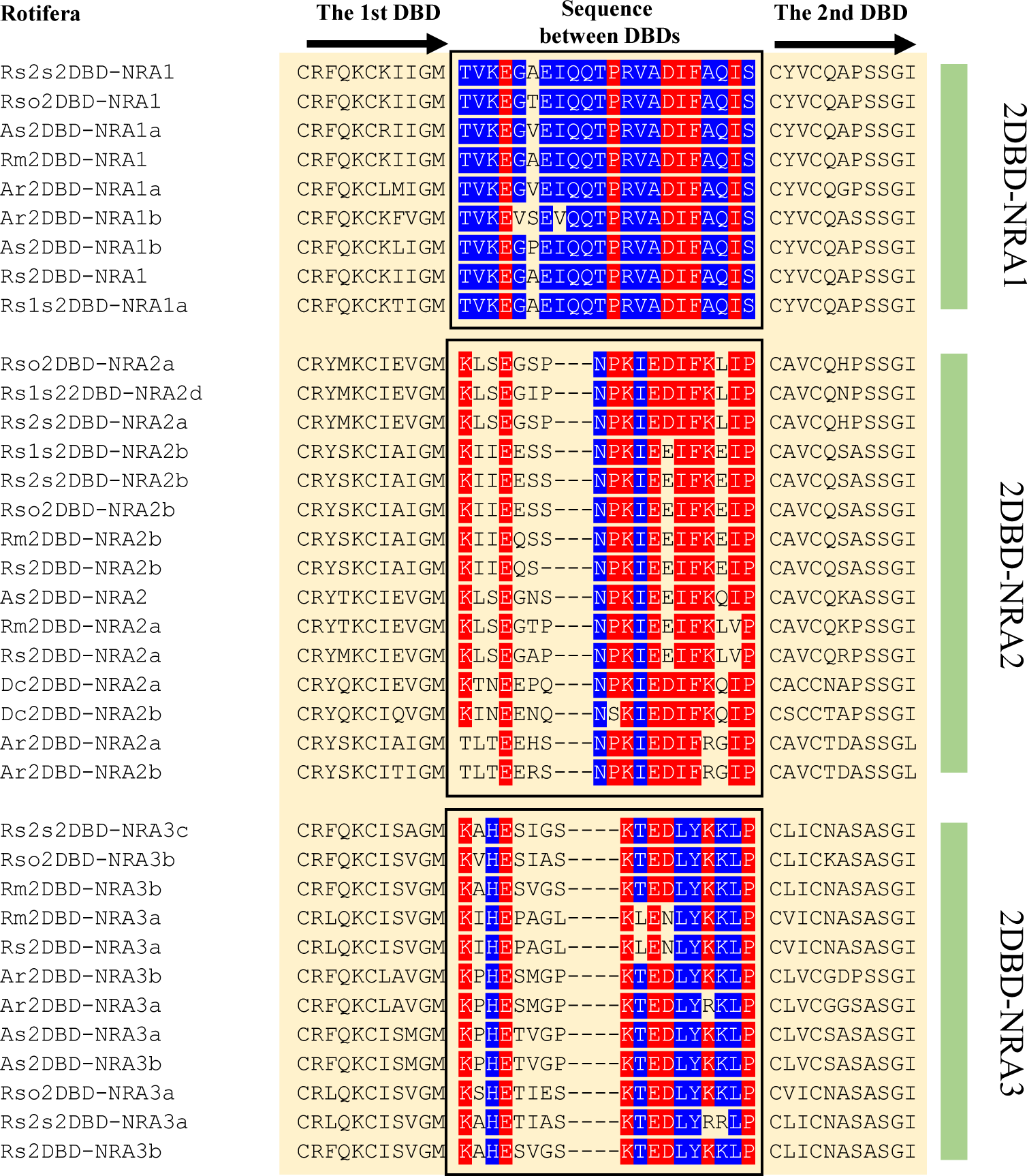
Amino acid sequence between the first and second DBDs in Rotifera 2DBD-NRs. Ar: *Adineta ricciae*, As: *Adineta steineri, Rotaria magnacalcarata*, Dc: *Didymodactylos carnosus*, Rm: *Rotaria magnacalcarata*, Rs: *Rotaria socialis*, Rs1s: *Rotaria sp. Silwood1*, Rs2s: *Rotaria sp. Silwood2,* Rso: *Rotaria sordida*, Rs1s: *Rotaria sp. Silwood1*. Blue highlighted letters indicate the conserved amino acid residues in each subgroup and red highlighted letters indicate the conserved amino acid residues among subgroups.

**Figure 6.**
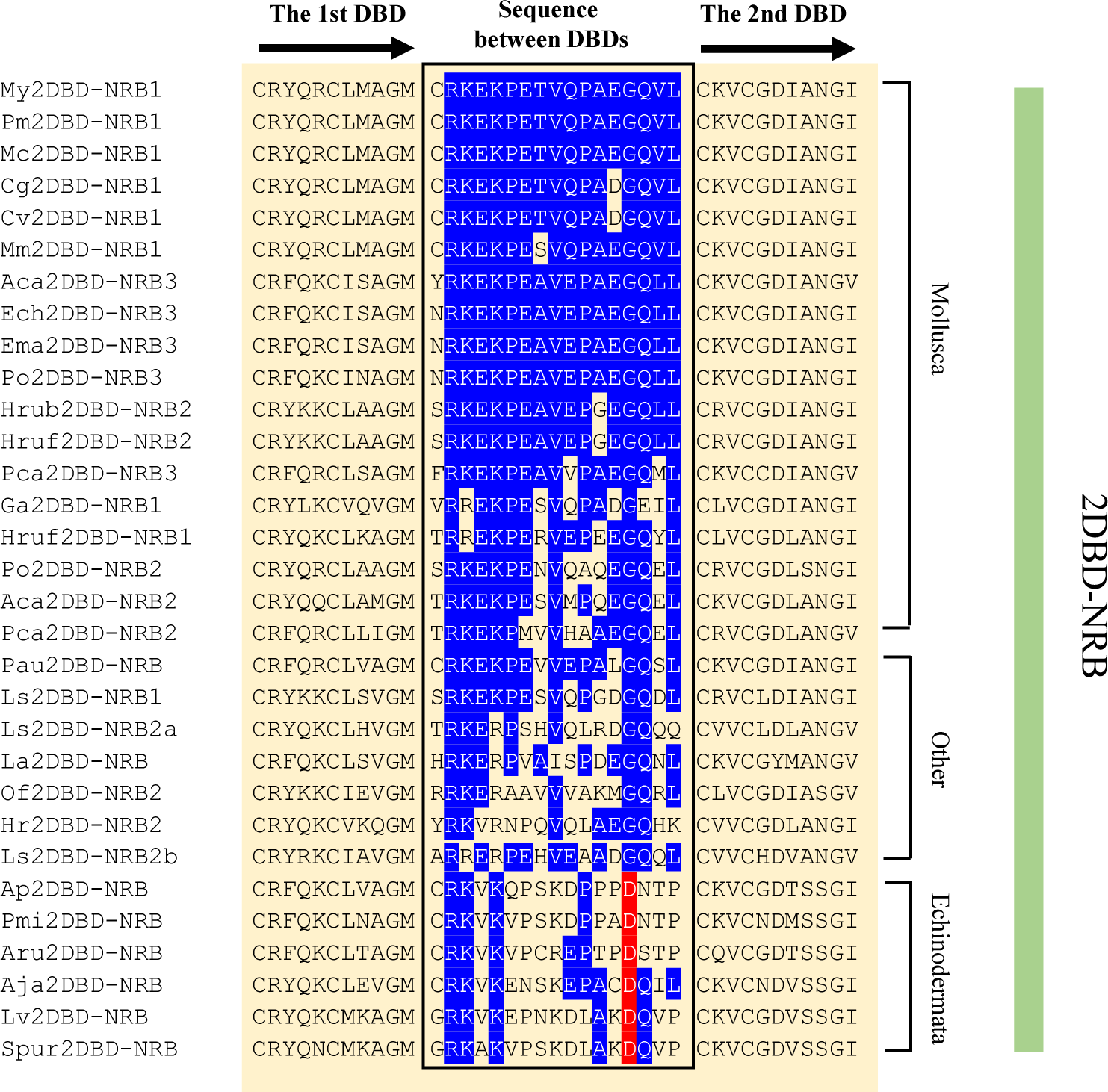
Amino acid sequence between the first and second DBDs in 2DBD-NRBs. Aca: *Aplysia californica*, Aj: *Anneissia japonica*, Ap: *Acanthaster planci*, Aru: *Asterias rubens*, Cg: *Crassostrea gigas*, Cv: *Crassostrea virginica*, Ech: *Elysia chlorotica*, Ema: *Elysia marginata*, Hr: *Helobdella robusta*, Hrub: *Haliotis rubra*, Hruf: *Haliotis rufescens*, La: *Lingula anatina*, Ls: *Lamellibrachia satsuma*, Lv: *Lytechinus variegatus*, Mco: *Mytilus coruscus*, Mm: *Mercenaria mercenaria*, My: *Mizuhopecten yessoensis*, Ofu: *Owenia fusiformis*, Pa: *Phoronis australis*, Pca: *Pomacea canaliculata*, Pm: *Pecten maximus*, Pmi: *Patiria miniata*, Po: *Plakobranchus ocellatus*, Spur: *Strongylocentrotus purpuratus*. Blue highlighted letters indicate the conserved amino acid residues in members of 2DBD-NRB group and red highlighted letters indicate the conserved amino acid residues in Echinodermata 2DBD-NRBs.

**4) The C-terminal Extension (CTE).** The C-terminal Extension (CTE) of DBD is important for DNA sequence recognition and binding. In 1992, Wilson et al. identified two boxes in CTE of human NGFI-B (hNR4A1). One box, termed T-Box, consisted of 12 amino acids that determined binding to tandem repeats of the half-site. The adjacent C-terminal seven amino acids, termed A-box, was required for recognition of the DNA binding element [35]. In 1998, Zhao et al. identified a subunit, termed Grip Box (G-box), in the CTE of human RevErbA-α. G-box formed a significant minor groove in the DNA binding surface [36]. They further showed that the G-box sequence was conserved in different orphan receptors despite the length of the pre-Grip sequences was different. The consensus G-box sequence they identified as RXGRZP (where X is a F, R, or G and Z usually contains a hydrophobic side chain). In this study, amino sequence alignment showed that all 2DBD-NRs contain a conserved G-box with the consensus sequence of RXGRQ(P/S) in 2DBD-NRAs, KXGR(P/H)X in 2DBD-NRBs and RDRRGP in Nematoda *A. avenae* 2DBD-NRCs (Fig. 7).

**Figure 7.**
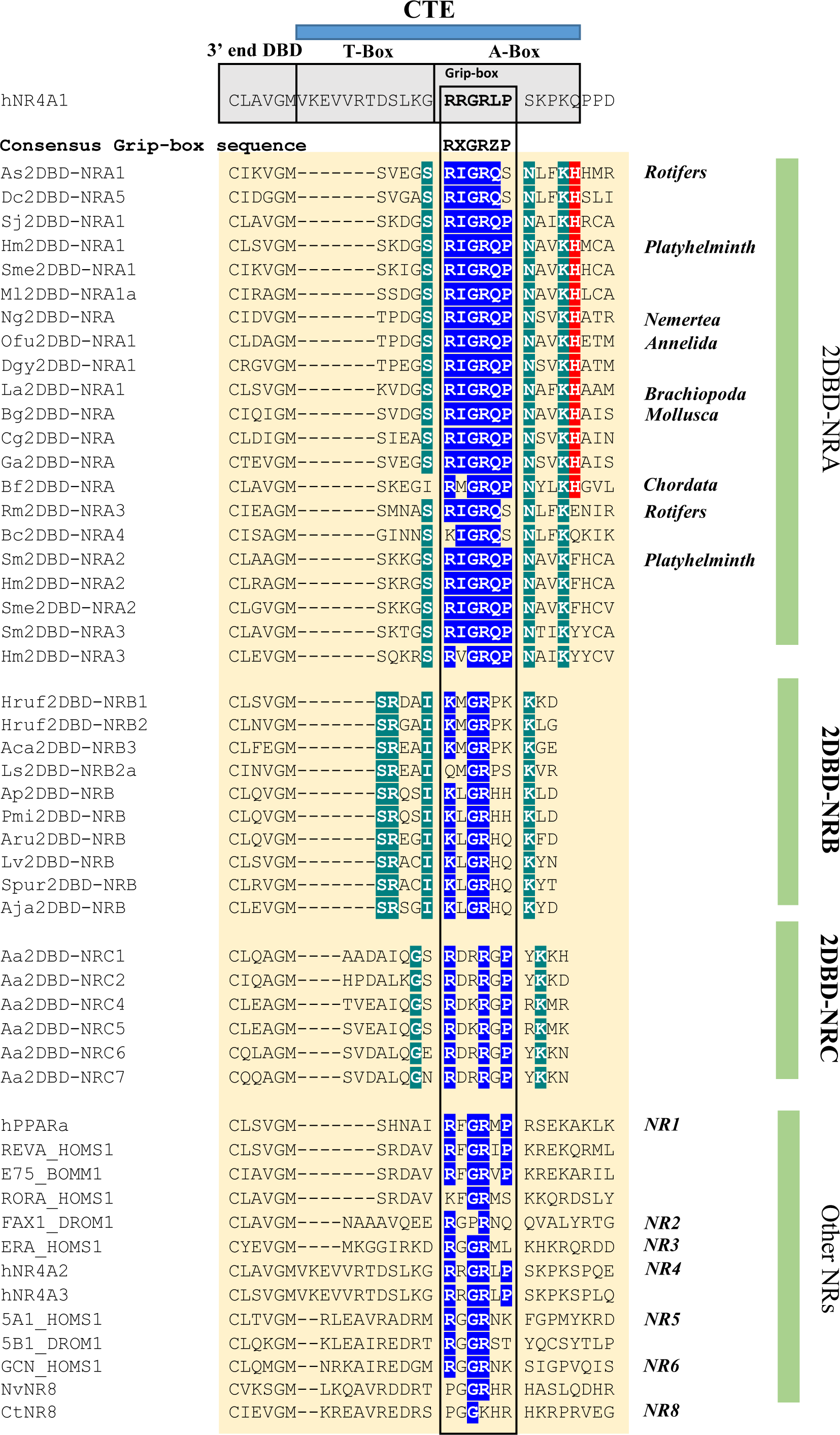
Amino acid sequence alignment of the C-terminal Extension (CTE) of 2DBD-NRs. Amino acid sequence alignment show conserved G-box sequence in 2DBD-NRs. The deep green highlighted letters indicate 2DBD-NR gene specific amino acid residues in CTE, the blue highlighted letters indicate conserved amino acid residues in the G-box of 2DBD-NRs and other NRs, the red highlighted letters indicate conserved amino acid residues amino acid (H) after the G-box. They are ancient signal amino acids among 2DBD-NRAs.

Using the length of the pre-Grip sequences and the sequence of the G-box, Zhao et al. divided NRs into three types (type-I, type-II, and type-III) [36]. Sequence alignment shows that there are five amino acid pre-Grip sequences that are the same as the type-I NRs as defined by Zhao et al [36]. Type I NRs include NRs that bind DNA as a monomer and homodimer (Rev-erb, E75 and Coup-TF), and heterodimer with RXR (PPAR). Our previous study showed that *S. mansoni* 2DBD-NRα (Sm2DBD-NRA2, in this study) could form a homodimer, but could not form a heterodimer with RXR [8]. The different conserved G-box sequence among 2DBD-NRA, 2DBD-NRB and 2DBD-NRC may represent the different DNA binding ability of these NRs.

Amino acid sequence alignment shows that the fifth amino acid (H) after the G-box are conserved in most 2DBD-NRAs, but not in Rotifer 2DBD-NRA3s and 2DBD-NRA4s, and not in Platyhelminth 2DBD-NRA2s and 2DBD-NRA3s. This suggests that the fifth amino acid residue (H) after the G-box are ancient signal amino acids among 2DBD-NRAs. The G-box sequence is also conserved in human NGFI-B that is localized in the 5’ end of the A-box of NGFI-B (Fig. 7), the pre-Grip sequences represents the T-box of NGFI-B which contains 12 residues [35]. The pre-Grip sequences in 2DBD-NR only contains five amino acids in 2DBD-NRA and 2DBD-NRBs, and eight amino acids in 2DBD-NRC, this data suggests that no conserved T-box is present in 2DBD-NRs.

**5) LBD.** LBD of NRs contain 12 helices and has two conserved regions, one region is known as “signature”, it contains 34 amino acid residues between the C terminus of H3 and the middle of H5. Within this region, a motif containing 20 amino acid residues was defined as an LBD specific signature (Ti) for the NR superfamily, it’s consensus sequence is ((F,W,Y)(A,S,I) (K,R,E,G)xxxx(F,L)xx(L,V,I)xxx(D,S)(Q,K)xx(L,V)(L,I,F)) [37]. In the C-terminus of LBD, there is an inducible transcription activation function TAF-2 (AF2) [38, 39]. The amino sequence of AF-2 region was conserved among many nuclear hormone receptors, the common consensus AF2-AD core structure is ФФxEФФ, where Ф denotes a hydrophobic residue [40–42]. Amino acid sequence alignment of 2DBD-NRs shows that Ti is conserved in 2DBD-NR (Fig. 8 and Supplemental material 3). AF2-AD core sequence is identified in most of 2DBD-NRs, but it is missing in nematode *Aphelenchus avenae* 2DBD-NRs. Sequence alignment shows that AF2-AD core sequence is conserved among 2DBD-NRs with ФФx(E,Q,R)ФФ in 2DBD-NRAs, ФФx(E,K)Фh (where h denotes a Hydrophilic residue) in 2DBD-NRBs and xxФФФФ in *Caenorhabditis brenneri* 2DBD-NRCs (Figure 8 and Supplemental material 4).

**Figure 8.**
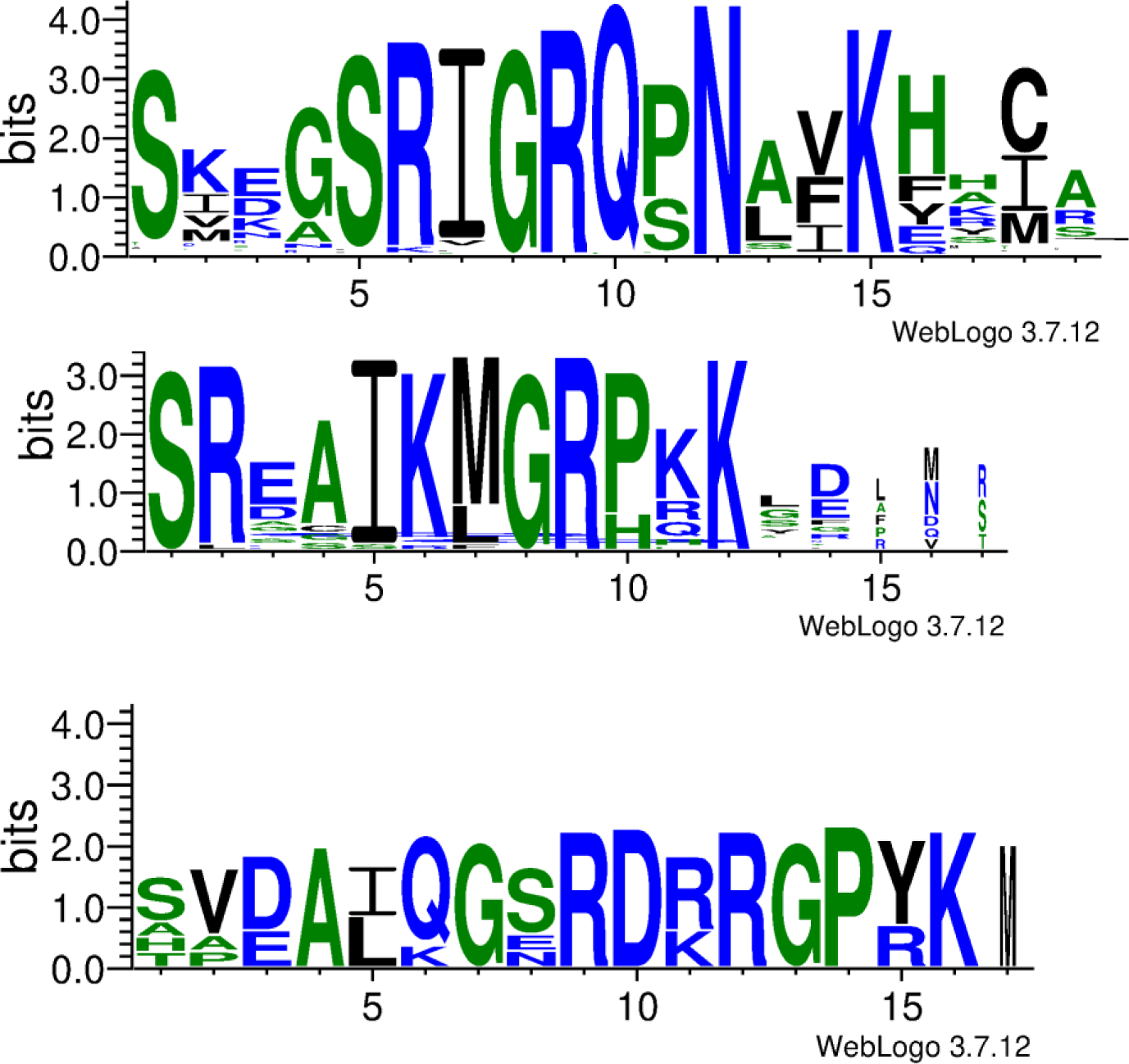
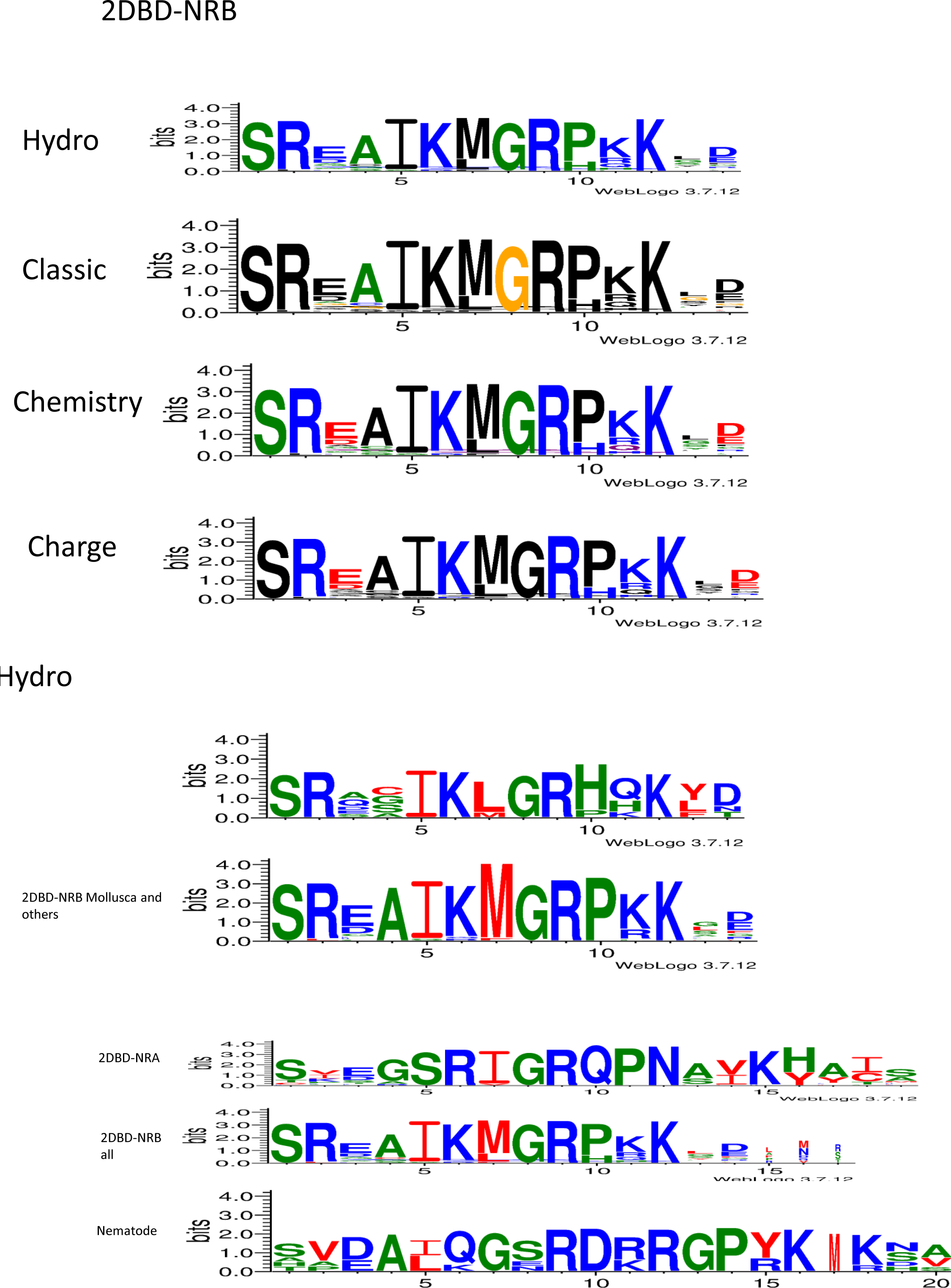
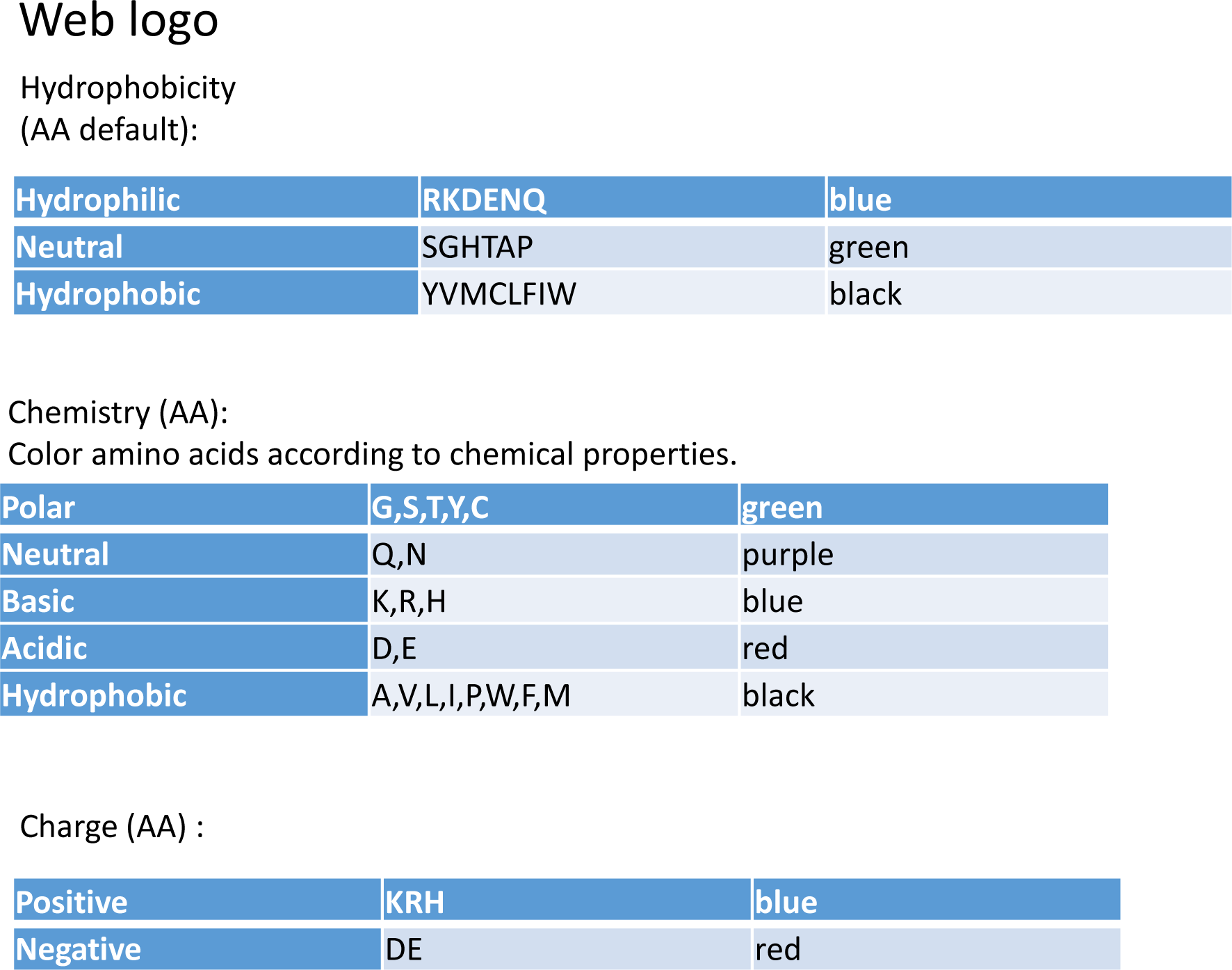
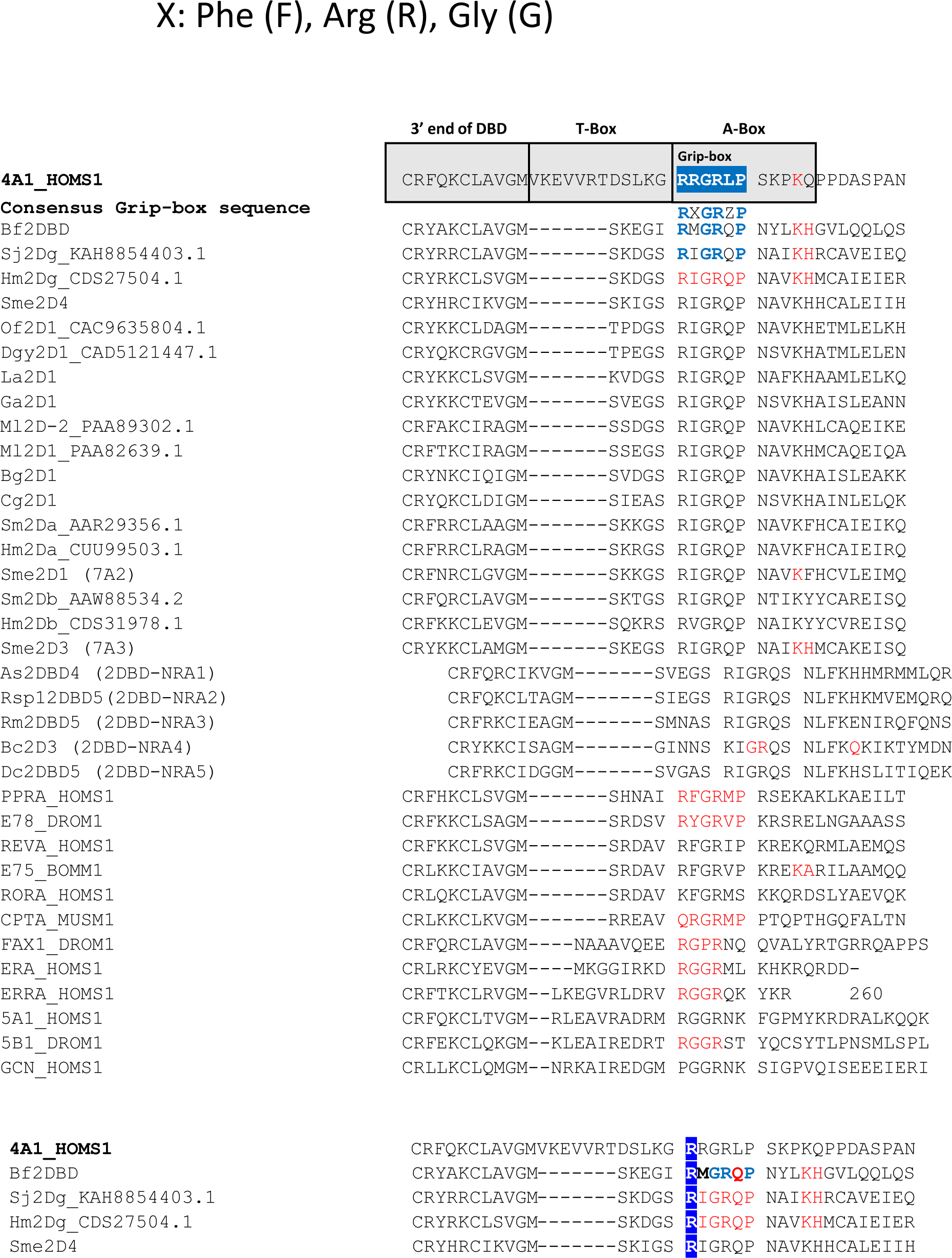
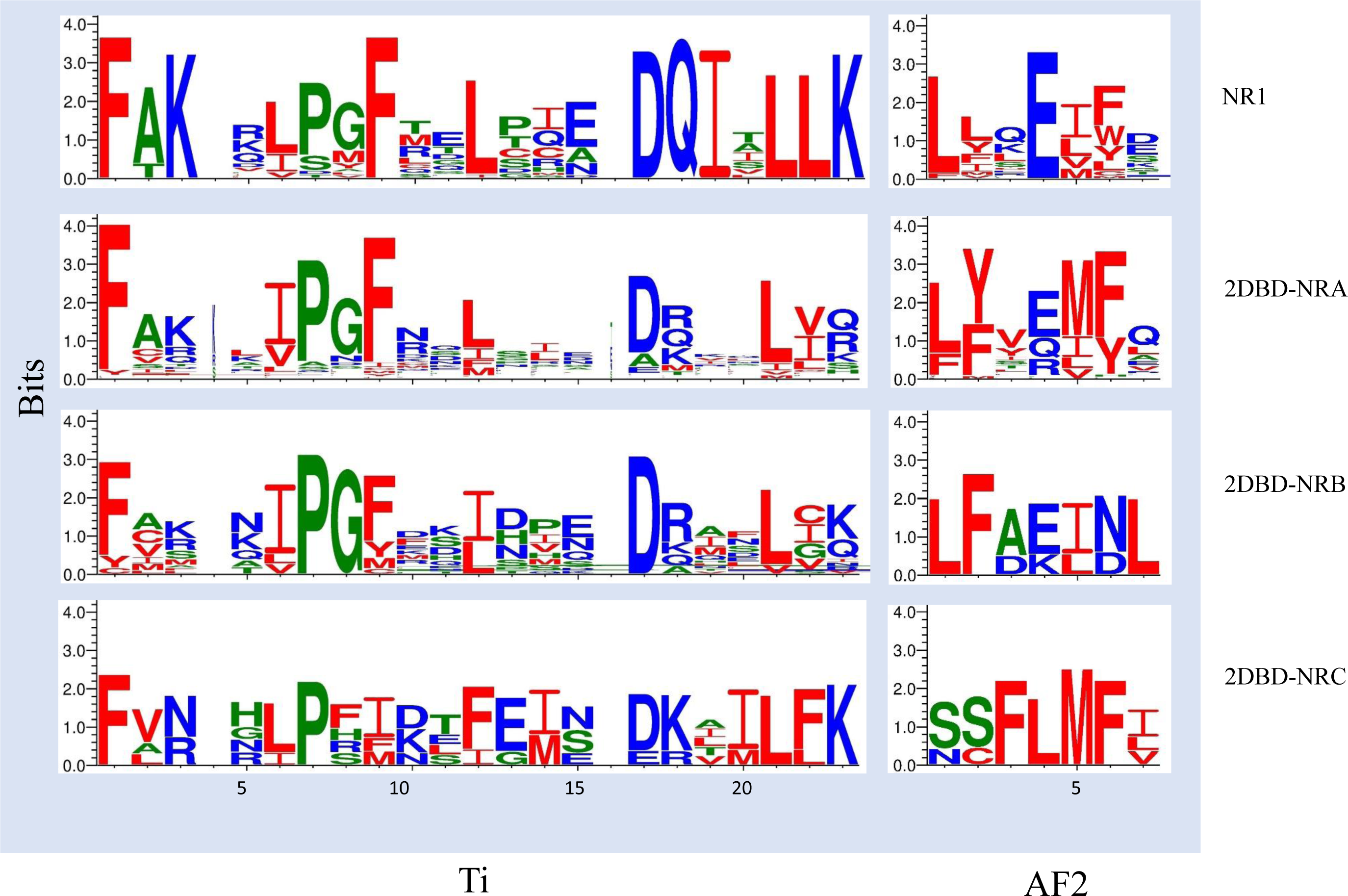
Conserved amino acid sequence of LBD specific signature (Ti) and activation function TAF-2 (AF2) in 2DBD-NRs. Sequence logo shows that both Ti and AF2 are conserved in 2DBD-NRs, and the pattern is similar to that of NR subfamily 1 (NR1). The sequence used for generation of the logo see supplemental material 3 and 4. Blue letter indicates a hydrophilic residue (RKDENQ), green letter indicates a neutral residue (SGHTAP) and red letter indicates a Hydrophobic residue (YVMCLFIW). [needs work, label figure 8]

### 4. A proposed nomenclature for 2DBD-NRs

A nomenclature for the NR superfamily was proposed, typical NRs (with both DBD and LBD) include 6 subfamilies (I-VI), and the NRs that are missing either DBD or LBD are placed in subfamily 0, irrespective of their evolutionary origin [43]. Due to the later finding of the 2DBD-NRs, they are not included in this classification system yet. According to our phylogenetic analysis of 2DBD-NRs, we propose a nomenclature for 2DBD-NRs following the rule of Nomenclature System for Nuclear Receptors [43]:

**1) NR subfamilies are designated by Arabic numerals (NR7, 2DBD-NR)**

Previously, we suggested placing 2DBD-NRs in a new NR subfamily (NR subfamily 7, NR7) when we discovered the 2DBD-NRs [6, 8] and this was accepted by some other studies [10, 44, 45]. However, NR7 was recently used to describe NRs [46] previously known as NR8 members [45] and amphioxus NR1Hs [27, 47]. To avoid confusion of 2DBD-NRs with other NRs named as NR7, our proposal keeps 2DBD-NRs as a subfamily instead of using NR7.

**2) Groups are designated by capital letters (2DBD-NRA, 2DBD-NRB and 2DBD-NRC)**

Phylogenetic analysis shows that 2DBD-NRs contain three groups: 2DBD-NRA, 2DBD-NRB and 2DBD-NRC (Fig. 2). Both 2DBD-NRA and 2DBD-NRB groups contain members from protostome Spiralia and deuterostomes, while 2DBD-NRC only contains members from nematode.

**3) Individual genes are designated by Arabic numerals**

For example, three 2DBD-NRs are identified in Mollusca *Aplysia californica*. One of them clustered in 2DBD-NRA group and is defined as *A. californica* 2DBD-NRA gene (Ac2DBD-NRA); two of *A. californica* 2DBD-NRs are clustered in 2DBD-NRB group, one is in Mollusca 2DBD-NRB2 subgroup 2, and it is defined as *A. californica* 2DBD-NRB2 (Ac2DBD-NRB2). The other one is clustered in Mollusca mono-phylogenetic 3 subgroup and it is defined as *A. californica* 2DBD-NRB3 (Ac2DBD-NRB3) (Fig. 2).

**4) A lowercase letter is added at the end of the gene to designate variants**

In Rotifera, 2DBD-NR genes underwent more rounds of gene duplications and gave birth to different gene variations, thus, a lowercase letter is added at the end of the gene to designate variants. For example, two variations of *Brachionus calyciflorus* 2DBD-NRs are clustered in Rotifera 2DBD-NRA1 subgroup, thus, they are defined as *B. calyciflorus* 2DBD-NRA1a (Bc2DBD-NRA1a) and *B. calyciflorus* 2DBD-NRA1b (2DBD-NRA1b), respectively. All identified 2DBD-NRs, their name in this nomenclature and their GenBank Accession number are listed in Table 6.

**Table 6.**
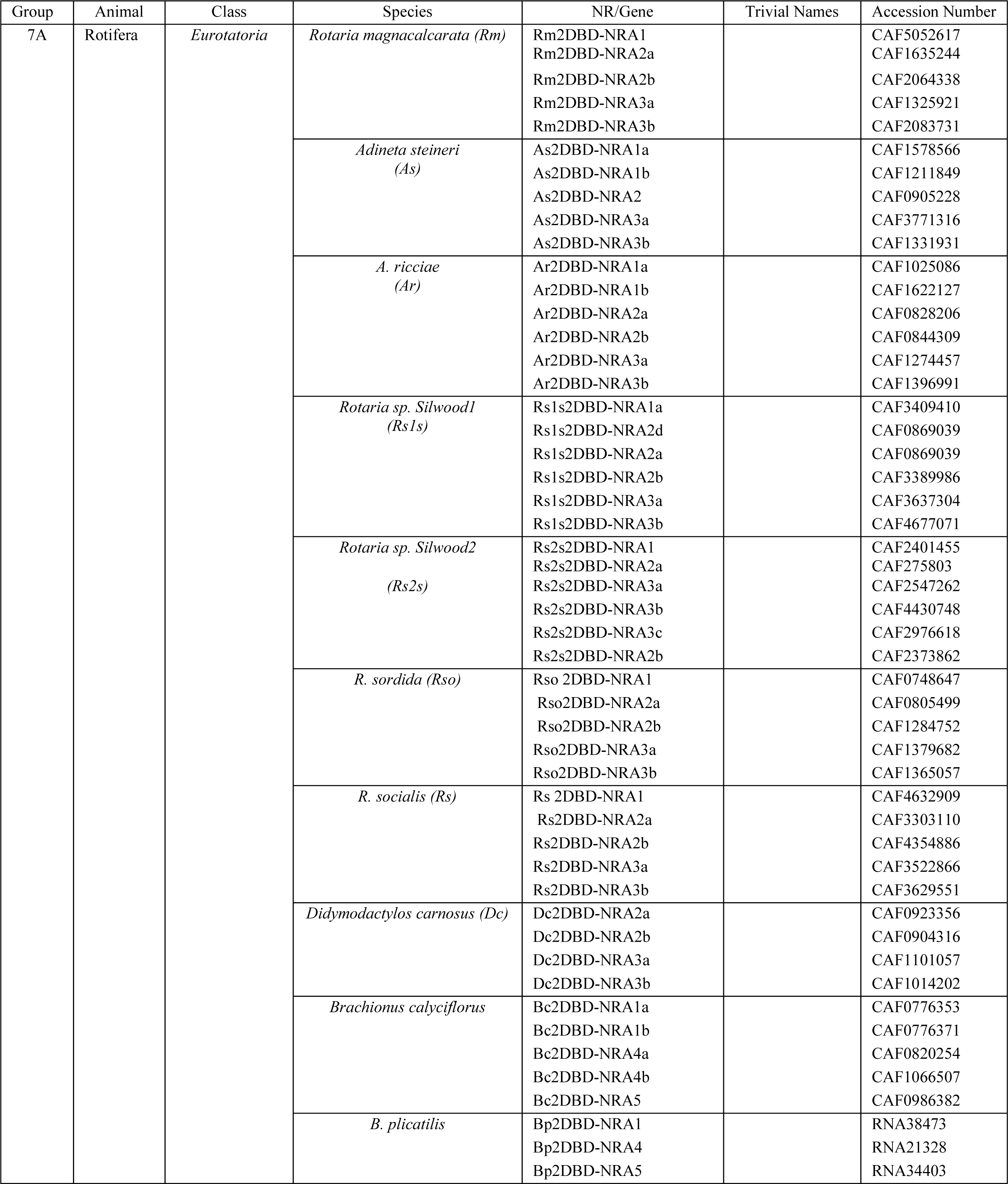
A Proposed Nomenclature for 2DBD-NRs.

| Group | Animal | Class | Species | NR/Gene | Trivial Names | Accession Number |
| --- | --- | --- | --- | --- | --- | --- |
| 7A | Rotifera | *Eurotatoria* | *Rotaria magnacalcarata (Rm)* | Rm2DBD-NRA1<br>Rm2DBD-NRA2a<br>Rm2DBD-NRA2b<br>Rm2DBD-NRA3a<br>Rm2DBD-NRA3b |  | CAF5052617<br>CAF1635244<br>CAF2064338<br>CAF1325921<br>CAF2083731 |
|  |  |  | *Adineta steineri (As)* | As2DBD-NRA1a<br>As2DBD-NRA1b<br>As2DBD-NRA2<br>As2DBD-NRA3a<br>As2DBD-NRA3b |  | CAF1578566<br>CAF1211849<br>CAF0905228<br>CAF3771316<br>CAF1331931 |
|  |  |  | *A. ricciae (Ar)* | Ar2DBD-NRA1a<br>Ar2DBD-NRA1b<br>Ar2DBD-NRA2a<br>Ar2DBD-NRA2b<br>Ar2DBD-NRA3a<br>Ar2DBD-NRA3b |  | CAF1025086<br>CAF1622127<br>CAF0828206<br>CAF0844309<br>CAF1274457<br>CAF1396991 |
|  |  |  | *Rotaria sp. Silwood1 (Rs1s)* | Rs1s2DBD-NRA1a<br>Rs1s2DBD-NRA2d<br>Rs1s2DBD-NRA2a<br>Rs1s2DBD-NRA2b<br>Rs1s2DBD-NRA3a<br>Rs1s2DBD-NRA3b |  | CAF3409410<br>CAF0869039<br>CAF0869039<br>CAF3389986<br>CAF3637304<br>CAF4677071 |
|  |  |  | *Rotaria sp. Silwood2 (Rs2s)* | Rs2s2DBD-NRA1<br>Rs2s2DBD-NRA2a<br>Rs2s2DBD-NRA3a<br>Rs2s2DBD-NRA3b<br>Rs2s2DBD-NRA3c<br>Rs2s2DBD-NRA2b |  | CAF2401455<br>CAF275803<br>CAF2547262<br>CAF4430748<br>CAF2976618<br>CAF2373862 |
|  |  |  | *R. sordida (Rso)* | Rso 2DBD-NRA1<br>Rso2DBD-NRA2a<br>Rso2DBD-NRA2b<br>Rso2DBD-NRA3a<br>Rso2DBD-NRA3b |  | CAF0748647<br>CAF0805499<br>CAF1284752<br>CAF1379682<br>CAF1365057 |
|  |  |  | *R. socialis (Rs)* | Rs 2DBD-NRA1<br>Rs2DBD-NRA2a<br>Rs2DBD-NRA2b<br>Rs2DBD-NRA3a<br>Rs2DBD-NRA3b |  | CAF4632909<br>CAF3303110<br>CAF4354886<br>CAF3522866<br>CAF3629551 |
|  |  |  | *Didymodactylos carnosus (Dc)* | Dc2DBD-NRA2a<br>Dc2DBD-NRA2b<br>Dc2DBD-NRA3a<br>Dc2DBD-NRA3b |  | CAF0923356<br>CAF0904316<br>CAF1101057<br>CAF1014202 |
|  |  |  | *Brachionus calyciflorus* | Bc2DBD-NRA1a<br>Bc2DBD-NRA1b<br>Bc2DBD-NRA4a<br>Bc2DBD-NRA4b<br>Bc2DBD-NRA5 |  | CAF0776353<br>CAF0776371<br>CAF0820254<br>CAF1066507<br>CAF0986382 |
|  |  |  | *B. plicatilis* | Bp2DBD-NRA1<br>Bp2DBD-NRA4<br>Bp2DBD-NRA5 |  | RNA38473<br>RNA21328<br>RNA34403 |

## CONCLUSION

In this study, 2DBD-NRs were identified in both protostomes and deuterostomes. Phylogenetic analysis shows that 2DBD-NRs consist of three groups, two groups are present in both protostomes and deuterostomes and the members of the other group is only found in Nematodes. Members of 2DBD-NRA and 2DBD-NRB are identified in both protostomes and deuterostomes, this result suggests that at least two 2DBD-NR genes were present in a common ancestor of the protostomes and deuterostomes. Phylogenetic analysis shows that 2DBD-NRs underwent gene duplication after the split of the different animal phyla. Thus, most of 2DBD-NR genes in a certain animal phyla are paralogues, rather than orthologues, of that in another animal phylum. 2DBD-NR gene losses occurred in different animal phyla, for example, 2DBD-NRA was missing in Phoronida and Echinodermata, and 2DBD-NRB was not identified in Nemertea, Rotifera, Platyhelminthes and Chordata. Sequence analysis shows that 2DBD-NRs possess highly conserved regions similar to that of typical NRs. The different P-P module (amino acid sequence of P-box in the first DBD and the second DBD) in different 2DBD-NR groups may affect their DBD binding abilities. Since very few studies have been carried out about 2DBD-NRs, little is known about their function. This study demonstrates that 2DBD-NR genes are widely distributed in both protostomes and deuterostomes, their role in regulation of the animal development awaits to be revealed.

