## Supplemental Material 1 for "Updated knowledge and a proposed nomenclature for nuclear receptors with two DNA Binding Domains (2DBD-NRs)"

Supplemental material 1. P-box sequence in 2DBD-NRs

| **2DBD-NR** | **P-box sequence** | |
|---|---|---|
| | **The first DBD** | **The second DBD** |
| **2DBD-NRA** | **EACKK** | **EGCKG** |
| Rotifera 2DBDs | | |
| Rotifera 2DBD-NRA3 group | EACKK | EACKG |
| Ar2DBD-NRA2b | EACKK | ESCKG |
| Hr2DBD-NRA3b | EACKK | ERCKG |
| Dc2DBD-NRA3b | EACKK | EACKC |
| **2DBD-NRB** | **LPCKS** | **EGCKK** |
| Ct2DBD-NRB2 | IPCKA | EGCKK |
| Hr2DBD-NRB2, Ls2DBD-NRB2a | VPCKT | EGCKK |
| Ls2DBD-NRB2b | LPCKT | EGCKK |
| La2DBD-NRB | LACKS | EGCKK |
| Of2DBD | WTCKT | LGCTK |
| **Echinodermata 2DBD-NRB** | **EACKS** | **EGCKG** |
| **2DBD-NRC** | | |
| Cb2DBD-NRC | DSCKC | DSCKW |
| Cr2DBD-NRC1 | DNCRA | GSCRM |
| Cr2DBD-NRC2 | DGCRT | EGCRV |
| Cr2DBD-NRC3 | EGCRS | EGCKG |
| Cr2DBD-NRC4 | EGCKK | EACKT |
| Aa2DBD-NRC1 | TGCKT | LGCKA |
| Aa2DBD-NRC2 | AGCRI | FGCKT |
| Aa2DBD-NRC3, Aa2DBD-NRC4 | NGCKT | HGCKA |
| Aa2DBD-NRC5 | NGCKT | HGCKS |
| Aa2DBD-NRC6 | TACWM | LGCKG |
| Aa2DBD-NRC7 | TACWM | HACKT |
