## Supplemental material 2 for "Updated knowledge and a proposed nomenclature for nuclear receptors with two DNA Binding Domains (2DBD-NRs)"

### Supplemental material 2. Sequence alignment of both DBDs of 2DBD-NRs

```
                                              The 1st DBD                                                                   The 2nd DBD
                 ________________________________________________________________________________________________________________________________________________________
                                     P-box                    D-Box                                                  P-box                     D-Box
                 CxxCxxx xxxxxxxxxxC xxC                      CxxxxxC xxxxxxxxxCxxC   CxxCxxxxxxxxxxxxxC xxC                       CxxxxxC    xxxxxxxxxCxxC
Ct2DBD-NRA1      CQVCGDV-AAGFHCGAYVC EACKK FFVRCLKQDA------FV CPRNNQC DITKDTRTQCQRCRYRKCLALGM CSVCGAPSSGFHFGAITC EGCKG FFRRT-IK--ERDI-LKYQ-- CSREGTC    IITTATRNICKFCRYQKCLDAGM
Ofu2DBD-NRA1     CQVCNDM-AAGFHCGAYVC EACKK FFLRCTKQD-------YM CGKQGSC IITRETRTQCQSCRYEKCLFLGM CRVCGARSSGYHFGAITC EGCKG FFRRT-IK--ERTA-DQYL-- CSRVGNC    EITLESRNSCKYCRYKKCLDAGM
Dgy2DBD-NRA1     CQVCNDV-AAGFHCGAYVC EACKK FFVRCQKQQ-------FA CQRNRDC VVTKETRTQCQSCRYQKCIQLGM CKVCGAPSSGYHFGAITC EGCKG FFRRT-VR--ERDS-CKYV-- CARNGRC    TITTATRNMCKFCRYQKCRGVGM
Ls2DBD-NRA1      CQVCNDI-AAGFHCGAYVC EACKK FFVRCLKQDT------FM CPRKKDC DITKETRTQCQYCRYQKCLAIGM CKVCGAPSSGYHFGAITC EGCKG FFRRT-VK--ERDS-LKYQ-- CSHFGNC    QITTQTRNVCKFCRYQKCLEVGM
Hr2DBD-NRA2      CQVCGDV-AAGFHCGAYVC EACKK FFLRCLKTGRGL---MFL CPKNSNC DITKETRSHCQRCRYQRCIALSM CKVCKAPSSGFHFGAITC EGCKG FFRRM-LK--EQVH-RKFI-- CSNDERC    NIDSANRTTCKSCRFNKCIKIGM
Ct2DBD-NRA2      CQVCGDI-AAGFHCGAYVC EACKK FYLRCLKSQANT---NFV CPKNSTC VVTKETRSHCQYCRFNKCIQLGM CKVCTAPSSGFHFGAITC EGCKG FFRRM-LK--EQLH-RKFQ-- CSGNDQC    DITSATRTTCKACRFSKCISVGM
Hr2DBD-NRA3c     CDVCADV-SSGFYCGAFVC EACKK FFVRCLQMKEKFR---AV CTKDGKC VINKTTRTQCSKCRYEKCISLNM CKVCGAPSSGFHFGVTTC EGCKG FYRRC-VK--KYDT-AKFH-- CTNDNNC    DVTWKTRNVCKFCRYQKCQDVGM
Hr2DBD-NRA3b     CQICGDS-AAGFYCGAYVC EACKK FFIRSLRSKISRDF--NP CPSDGQC HVTKETRTQCPQCRYNKCLSFNM CKVCDFPSSGFHYGAITC ERCKG FFRRC-IL--DRNI-ELLG-- CM????C    RLLKKKRNMCKRCRYLKCLAVGM
Hr2DBD-NRA3a     CQVCTDP-AAAFYCGAQVC EACKK FFIRSWKNSTENN---YV CLQDRKC VLTKESRKHCAYCRYDRCLQLKM CRICGAPSSGFHFGVITC EGCKG FFRRR-CH--DNRF-DKFK-- CNENNCC    VISAANRSMCRACRLRKCLDSGM
Em2DBD-NRA1      CDICGDV-SAGFHCNAYVC EACKK FFIRSSKNENY-----YS CSKVNAC EVNKDTRTHCQRCRFYKCLRIGM CRVCGAQSSGFHFGAITC EGCKG FFRRT-IS--ERDN-QRYT-- CRNGGTC    LITLATRNNCKSCRYRKCLSVGM
Sm2DBD-NRA1      CDICGDV-AAGFHCNAYVC EACKK FFIRSSKGENF-----YT CTKSNTC EINKDTRTHCQRCRYQKCIRLGM CRVCGAKSSGFHFGAITC EGCKG FFRRT-IN--ERES-QRYT-- CRNGGNC    AVTGATRNNCKSCRYRRCLAVGM
Px2DBD-NRA1      CDICGDV-SAGYHCNAYVC EACKK FFIRSSKGRSY-----YT CTKASNC DIKKDTRTHCQYCRFQKCLHIGM CRVCGAKSSGFHFGAITC EGCKG FFRRT-VN--ERDG-HRYT-- CRNGGNC    SVTGSTRNNCKSCRYQRCLLVGM
Sme2DBD-NRA1     CDVCGDV-SAGFHCSAFVC EACKK FFIRSSKGDSY-----YS CTKNNNC EIVKDTRTHCQYCRFQKCLHLGM CRVCGAQSSGFHFGAITC EGCKG FFRRT-IN--EREN-QKYT-- CRNGGNC    VINLATRNNCKSCRYKKCLAMGM
Ml2DBD-NRA1a     CDVCGDT-AAGFHCGAFVC EACKK FFIRSTRAGMQ-----YS CSKVGRC DVTKETRTHCQHCRYKKCLMLNM CRVCGASSSGFHFGAITC EGCKG FFRRT-IK--ERDA-GKYI-- CSKGGGC    EINQQSRNACKSCRFAKCIRAGM
Ml2DBD-NRA1b     CDVCGDT-AAGFHCGAFVC EACKK FFIRSSRAEQQ-----YA CSKSGSC DITKETRTHCQFCRYKKCLALRM CRVCGASSSGFHFGAITC EGCKG FFRRT-IK--ERDS-GKYV-- CSKGGGC    EINKSSRNACKSCRFTKCIRAGM
Sm2DBD-NRA2      CQVCGEL-AAGFHHGAYVC EACKK FFMRHSMADTKP---TNV CPTGGNC IVAKGSRGKCQICRYRKCLLVGM CRVCGGRSSGFHFGALTC EGCKG FFRRT-EG--SS---NSLV-- CVGQNAC    TITPRSRNACKSCRFRRCLAAGM
Px2DBD-NRA2      CQVCGEL-AAGFHHGAYVC EACKK FFMRHSLNDSKP---TNS CPSAGNC VVAKGSRGKCQQCRYRKCLDVGM CRVCGGRSSGFHFGALTC EGCKG FFRRT-EG--TS---QRLA-- CVGQDNC    AITPRSRNACKGCRFRRCLTAGM
Em2DBD-NRA2      CQICGEL-AAGFHHGAYVC EACKK FFMRHSLANGRS---TTT CPSGGNC VIAKSSRGRCQQCRYKKCLEVGM CRVCGGRSSGFHFGALTC EGCKG FFRRT-EE--SA---SRLV-- CVGKDNC    TITPRSRNVCKACRFRRCLRAGM
Sme2DBD-NRA2     CQVCRES-AAGFHHGAYVC EACKK FFMRHNLNSTKF---TIP CPTGGQC AQLKTGRIKCQSCRFKKCVSIGM CRVCGGKSSGFHFGALTC EGCKG FFRRT-ES--TK---QHLA-- CIGNNNC    CLTGSSRNLCKSCRFNRCLGVGM
Px2DBD-NRA3      CQVCGQP-AVGFHHRAYVC EACKK FFTRHASARLRSA-SSMA CPMGGGC RVEGPGRGKCPHCRFKKCLDLGM CRVCSGPSSGFHFGALTC EGCKG FFRRT-VL--SG---VHLE-- CLNNNDC    PVTPSNRNMCKACRFNRCLAVGM
Sm2DBD-NRA3      CQICGQP-AVGFHHRAYVC EACKK FFMRHTAARLRNS----I CPMGGRC RVEGPGRGKCPHCRYRKCLELGM CRVCSGPSSGFHFGALTC EGCKG FFRRT-VL--SN---VRLE-- CPGNNDC    PITPANRNMCKSCRFQRCLAVGM
Em2DBD-NRA3      CQVCGQP-SVGFHHRAYVC EACKK FFTRHLTNRIKKEKSSVV CPMGGNC KIEGPGRGKCPHCRFRKCLDLGM CRVCGGPSSGFHFGALTC EGCKG FFRRT-VH--SN---SIPQ-- CLGNQTC    RITPSNRNMCKSCRFKKCLEVGM
Sme2DBD-NRA3     CQVCNCL-AAGFHHGAYVC EACKK FFKRYSTNQLNSK---VV CKK--NC DLEKIGRRKCSFCRYQKCLNVGM CRVCGGKSSGFHFGSLTC EGCKG FFRRA-EP--LK---DKLE-- CVGKLNC    VIDSSHKMPCKWCRYHRCIKVGM
Sme2DBD-NRA4     CQICTKN-SVGFHYGAYVC EACKK FFVRHASGVCRLY---GS CQEKGKC DLNIDGRGKCQHCRYQRCNEIGM CQVCGAKSSGLHFGCITC EGCKG FFRRM-IK--FK---GSLV-- CFNDNKC    KLDLKNRSTCKSCRMNRCLSVGM
Ng2DBD-NRA       CKVCGDM-SSGFHCGAYVC EACKK FFIRSMKHSS------YV CSKNKTC RITKESRTQCQYCRYQKCLSSGM CKVCGAPSSGFHFGAITC EGCKG FFRRT-IK--ERDS-YRYA-- CSKGGNC    SITLSTRNACKYCRFKQCIDVGM
La2DBD-NRA1      CQVCGDL-SSGFYCGAFVC EACKK FYLRSLK-QNSK----YS CLKKGQC QVSKETRTVCQYCRFRKCIAIGM CRVCGAPSSGFHFGAVTC EGCKG FFRRT-VR--ERD--NYYS-- CSKRQQC    SITETTRNICKYCRYKKCLSVGM
La2DBD-NRA2      CSVCGET-SSGFHCGAITC EACKK FFIRSCQGESNK----YV CTKNNNC EIMRATRTQCQRCRYEKCLAVGM CSVCKAPASGFHFGAVTC EGCKG FFRRS-VK--EKNGLDYY--- CEQGGNC    PITETTRNTCRYCRYKKCLDVGM
Bl2DBD-NRA_      CQVCGDV-SVGFHCGACVC EACKK FFIRSTKDGGDR--TKFK CSKQQKC DITKQTRSQCQYCRLQKCLAVGM CKVCGAESSGFHFGVDTC EGCKG FFRRS-QL--KTEY-ESYR-- CSKSKDC    AINVVTRNLCRYCRYAKCLTVGM
Bf2DBD-NRA       CQVCGDV-SVGFHCGACVC EACKK FFIRSTKDGGDR--TKFK CSKQQKC DITKQTRSQCQYCRLQKCLAVGM CKVCGAESSGFHFGVDTC EGCKG FFRRS-QL--KTEY-ESYR-- CSKSKDC    AINVVTRNLCRYCRYAKCLAVGM
Cv2DBD-NRA       CQVCGDK-AAGFYCGAFVC EACKK FFMRASRQQKIK----YT CLKAKKC RITKESRVQCQYCRFQKCLSLNM CQVCNAPSSGFHFGALTC EGCKG FFRRI-AK--EKGS-KPYR-- CSKSQNC    VVTTGTRNLCRACRYQKCLDIGM
Cg2DBD-NRA       CQVCGDK-AAGFYCGAFVC EACKK FFMRASRQQKVK----YT CLKAKNC KITKESRVQCQYCRFQKCLSLNM CQVCNAPSSGFHFGALTC EGCKG FFRRI-AK--EKGN-KQYR-- CSKSQNC    EVTTGTRNLCRACRYQKCLDIGM
Pm2DBD-NRA       CQVCGDK-AAGFYCGAFIC EACKK FFMRASKLEKIK----YV CLRVQKC VITKESRVQCQYCRFQKCLSLKM CRVCSAPSSGFHFGALTC EGCKG FFRRM-AK--EREC-QRYK-- CSKNGNC    DVNTVTRNLCKACRYRKCLESGM
My2DBD-NRA       CQVCGDK-AAGFYCGAFIC EACKK FFMRANKLEKLK----YV CLRVGRC VITKESRVQCQYCRYQKCVSLKM CRVCSAPSSGFHFGALTC EGCKG FFRRM-AK--EREC-QRYK-- CSKNGSC    EVNTITRNLCKACRYRKCVECGM
Dp2DBD-NRA       CQVCGDV-AAGFYCGAYIC EACKK FYMRAYKQDKLK----YV CLRDKKC VITKESRVQCQYCRFQKCVQLNM CRVCSAPSSGFHFGAITC EGCKG FFRRM-AK--EREA-EKYQ-- CNKSGDC    EINMVTRNLCKACRYRACIKAGM
Mm2DBD-NRA       CQVCGDM-AAGFYCGAFIC EACKK FYMRAYKQDKLK----YV CLRDKKC MITKESRVQCQYCRFQKCVTLNM CRVCSAPSSGFHFGAITC EGCKG FFRRM-AK--EREP-EKYQ-- CNKKGRC    EINMVTRNLCKACRYRSCREAGM
Mg2DBD-NRA       CQVCGDK-AAGFYCGAFAC EACKK FFMRSCRNDKTK----YV CLRDKKC QMTAESRVQCQYCRLQKCLSLNM CRVCGAPSSGFHFGALTC EGCKG FFRRM-AK--EREL-GKYR-- CSKTGSC    EINTSTRNLCKSCRYKKCVDAGM
Me2DBD-NRA       CQVCGDK-AAGFYCGAFAC EACKK FFMRSCRNDKTK----YV CLRDKKC QMTAESRVQCQYCRLQKCLSLNM CRVCGAPSSGFHFGALTC EGCKG FFRRM-AK--EREL-GKYR-- CSKTGSC    EINTSTRNLCKSCRYKKCVDAGM
Mc2DBD-NRA       CQVCGDK-AAGFYCGAFAC EACKK FFMRSCRNDKTK----YV CLRDKKC QMTAESRVQCQYCRLQKCLSLNM CRVCSAPSSGFHFGALTC EGCKG FFRRM-AK--EREL-GKYR-- CSKTGSC    EINTSTRNLCKSCRYKKCVDAGM
Ech2DBD-NRA 1    CQVCGDQ-AAGFYCGAYIC EACKK FFIRASKLKQMK----YV CLKQKNC NITKETRVHCQFCRYQKCLSLNM CRVCSAPSSGFHFGALTC EGCKG FFRRM-VN--EREP-GTYS-- CGRGGNC    EVNSMTRNMCKACRYAKCLQIGM
Ema2DBD-NRA      CQVCGDQ-AAGFYCGAYIC EACKK FFIRASKLKQMK----YI CLKQKNC KITKESRVHCQYCRYQKCLSLDM CRVCSAPSSGFHFGALTC EGCKG FFRRM-VN--EREP-GTYT-- CGKGGKC    EVNSMTRNMCKACRYAKCLMIGM
Po2DBD-NRA       CQVCGDH-AAGFYCGAYIC EACKK FFIRAAKLKQIK----YV CLKQKNC IITKESRVHCQYCRYQKCLSLNM CRVCSAPSSGFHFGALTC EGCKG FFRRM-VN--EREP-GTYK-- CGKGGNC    KVNSMTRNMCKACRYAKCLQIGM
Bg2DBD-NRA       CQVCGDL-AAGFYCGAYIC EACKK FFIRASKLERPR----YV CLRQKDC VITKESRVHCQYCRYQKCLQLNM CRVCAAPSSGFHFGALTC EGCKG FFRRM-VK--EREA-YNYK-- CSKSGTC    EVNAMTRNMCKACRYNKCIQIGM
Aca2DBD-NRA      CQVCDDL-AAGFYCGAYIC EACKK FFIRASKLECPR----YV CLRQKNC IITKESRVHCQFCRYQKCLQLNM CRVCSAPSSGFHFGALTC EGCKG FFRRM-VK--EREA-CAYK-- CSKDGNC    EVNAMTRNMCKACRYHKCLQIGM
Cu2DBD-NRA       CQVCEDA-AAGFYCGAYIC EACKK FFIRASKQNELR----YQ CPRSRSC PITRESRVHCQYCRYQKCLRLSM CRVCSAPSSGFHFGALTC EGCKG FFRRM-VK--ERDS-YSYK-- CGRNGHC    EINATTRNMCKACRYNKCLQIGM
Pca2DBD-NRA      CQVCGDI-AAGFYCGAFIC EACKK FFIRASKQEKHK----FV CLRQGLC KITKESRVQCQFCRYQKCLDLQM CRVCSAPSSGFHFGALTC EGCKG FFRRM-VK--EREP-GSYR-- CSAGGTC    EISAQTRNMCKACRYQKCVDIGM
Hruf2DBD-NRA     CQVCGDV-AAGFYCGAYIC EACKK FFVRASRQEKVK----FV CHKSEDC VITKETRVRCQHCRYQKCLTLHM CRVCGGASSGFHFGALTC EGCKG FFRRM-AK--EREW-HSYK-- CSKEDRC    EISMLTRNVCKACRYRKCLDAGM
Hrub2DBD-NRA     CQVCGDV-AAGFYCGAYIC EACKK FFVRASRQEKVK----FV CHKSEDC VITRETRVRCQHCRYQKCLSLDM CRVCGGSSSGFHFGALTC EGCKG FFRRM-AK--EREW-HSYK-- CSKDDKC    EISMLTRNVCKACRYRKCLEAGM
Ga2DBD-NRA       CQVCGDV-AAGFYCGAFIC EACKK FFVRTCKQDKVK----YA CHKNQCC KITKENRIRCQYCRYQKCLSVNM CRVCGGPSSGFHFGALTC EGCKG FFRRM-VK--ERSP-SMYQ-- CLKDNDC    NINMFTRNICKSCRYKKCTEVGM
Lg2DBD-NRA       CDVCGDK-AAGFYCGAFVC EACKK FFIRAAKQGEVK----YK CLRDGNC TITKVNRIQCQFCRYQKCVALNM CQVCGAQSSGFHFGALTC EGCKG FFRRM-AK--ERSS-NSYI-- CSKGNMC    LVSISTRNSCKACRYQKCILVGM
Rs1s2DBD-NRA2a   CQVCGEN-ASGWHCGSITC EACKK FFLRSVNEEYR-----YK CMKDKKC IITRTTRTQCQYCRYMKCIEVGM CAVCQNPSSGIHFGATTC EGCKG FFRRT-MK--ERTP-QRYK-- CPEKNNC    EVNTSTRSACRACRFQKCLTAGM
Rs2s2DBD-NRA2    CQVCGEN-ASGWHCGSITC EACKK FFLRSVNEEYR-----YK CMKDKKC IITRTTRTQCQYCRYMKCIEVGM CAVCQHPSSGIHFGATTC EGCKG FFRRT-MK--ERTP-QRYK-- CPEKNNC    EVNTSTRSACRACRFQKCLTAGM
Rso2DBD-NRA2a    CQVCGEN-ASGWHCGSITC EACKK FFLRSVNEEYR-----YR CMKDKKC IITRTTRTQCQYCRYMKCIEVGM CAVCQHPSSGIHFGATTC EGCKG FFRRT-MK--ERTP-QRYK-- CPEKNNC    EVNTSTRSACRACRFQKCLTAGM
As2DBD-NRA2      CQVCGEN-ASGWHCGSITC EACKK FFLRSVNDEYR-----YK CIKDKKC PITRTTRTQCQYCRYTKCIEVGM CAVCQKASSGIHFGATTC EGCKG FFRRT-IK--ERIP-QRYK-- CPEKNNC    EINYLTRSACRACRFQKCLTVGM
Ar2DBD-NRA2a     CQVCGEN-ASGWHCGSITC EACKK FFLRSVNDEYR-----YK CMKDKNC LITRTTRTQCQYCRYAKCIQVGM CAVCHHASSGIHFGATTC EGCKG FFRRT-MR--ERIP-PRYK-- CAENNTC    VIGYLTRNSCRACRFEKCLRAGM
Rs2DBD-NRA2a     CQVCGEN-ASGWHCGSITC EACKK FFLRSVNEEYR-----YK CIRDKKC IITRTTRTQCQYCRYMKCIEVGM CAVCQRPSSGIHFGATTC EGCKG FFRRT-IK--ERTP-QRYK-- CPENYNC    EVNSSTRSACRACRFQKCLAAGM
Rm2DBD-NRA2a     CQVCGEN-ASGWHCGSITC EACKK FFLRSVNEEYR-----YK CIRDKKC IITRTTRTQCQYCRYTKCIEVGM CAVCQKPSSGIHFGATTC EGCKG FFRRT-IK--ERTP-QRYK-- CPEDYNC    EVNSSTRSACRACRFQKCLAAGM
Rso2DBD-NRA2b    CQVCGEN-ASGWHCGSITC EACKK FFLRSVNGEYL-----YK CIRDKKC IITRTTRTQCQYCRYSKCIAIGM CAVCQSASSGIHFGATTC EGCKG FFRRM-IK--ERIP-QHYK-- CSERNNC    EVNALTRNMCRACRFRKCLMAGM
Rs2DBD-NRA2b     CQVCGEN-ASGWHCGSITC EACKK FFLRSVNGEYL-----YK CIRDKKC IITRATRTQCQCCRYSKCIAIGM CAVCQSASSGIHFGATTC EGCKG FFRRM-TK--ENTP-QNYK-- CSERNNC    EINALTRNMCRACRFRKCLMAGM
Rm2DBD-NRA2b     CQVCGEN-ASGWHCGSITC EACKK FFLRSVNGEYL-----YK CIRDKKC IITRTTRTQCQCCRYSKCIAIGM CAVCQSASSGIHFGATTC EGCKG FFRRM-TK--ENTP-QNYK-- CSERSNC    EINALTRNMCRACRFRKCLMAGM
Rs1s2DBD-NRA2b   CQVCGEN-ASGWHCGSITC EACKK FFLRSVNGEYL-----YK CIRDKKC IITRTTRTQCQYCRYSKCIAIGM CAVCQSASSGIHFGATTC EGCKG FFRRM-IK--ERIP-QHYK-- CSEKNNC    EVNAITRNMCRACRFRKCLVAGM
Rs1s2DBD-NRA2c   CQVCGEN-ASGWHCGSITC EACKK FFLRSVNGEYL-----YK CIRDKKC IIARTTRTQCQYCRYSKCIAIGM CAVCQSASSGIHFGATTC EGCKG FFRRM-IK--ERIP-QHYK-- CSEKNNC    EVNAITRNMCRACRFRKCLTAGM
Ar2DBD-NRA2b     CQVCGEN-ASGWHCGSITC EACKK FFLRSVNGEYL-----YR CIRDKKC PITRTTRTQCQFCRYSKCITIGM CAVCTDASSGLHFGVTTC ESCKG FFRRM-IK--ERIP-QHYK-- CLERNNC    EVNTSTRNLCRACRFQKCLSVGM
Dc2DBD-NRA2a     CQVCSEQ-SSGWHCGAVTC EACKK FFLRSVNEEHL-----YK CIRDKRC QVTRSTRTQCQYCRYQKCIEVGM CACCNAPSSGIHFGATTC EGCKG FFRRT-IK--ERTP-QRYK-- CLESNNC    EINTSTRNMCRACRFQKCVNAGM
Dc2DBD-NRA2b     CQVCSEQ-SSGWHCGAVTC EACKK FFLRSINEEYL-----YK CVRDKKC LITRSTRTQCQYCRYQKCIQVGM CSCCTAPSSGIHFGAMTC EGCKG FFRRT-MK--ERTS-QRYT-- CLESNNC    EINTSTRNICRACRFQKCINAGM
Ar2DBD-NRA3b     CQICGDL-SSGWHCGAITC EACKK FFLRSISTGDG-----YK CQRDFSC AITKRSRTQCQYCRFQKCLAVGM CLVCGDPSSGIHFGAVTC EACKG FFRRS-MR--ENAV-ERYR-- CAENNNC    EIRSGSKVTCRACRFRKCVQKGM
Ar2DBD-NRA3a     CQICGDL-SSGWHCGAITC EACKK FFLRSISTGDG-----YK CQRDFSC AITKRSRTQCQYCRFQKCLAVGM CLVCGGSASGIHFGAVTC EACKG FFRRS-IK--ENAP-DRYR-- CTENNNC    DIHATSKITCRACRFKKCVQAGM
As2DBD-NRA3a     CQICGDL-ASGWHCGAITC EACKK FFLRSISTGDG-----YK CQKDFSC SITKRSRTQCQYCRFQKCISMGM CLVCSASASGIHFGAVTC EACKG FFRRS-IK--ENAP-ERYR-- CAENNNC    EILSTSKITCRACRFKKCVDAGM
As2DBD-NRA3b     CQICGDL-ASGWHCGAITC EACKK FFLRSISTGDG-----YK CQKDFSC SITKRSRTQCQYCRFQKCISMGM CLVCSASASGIHFGAVTC EACKG FYRRS-IK--ENAA-ERYL-- CVENNSC    EILAPSKLSCRACRFKKCIHFGM
Dc2DBD-NRA3a     CQICCDT-ASGWHCGAITC EACKK FFLRSITTGDG-----YK CQRDQCC LINKRSRTQCQFCRFQKCLKAGM CLVCGAPASGIHFGAVTC EACKG FFRRS-IK--ENAP-DRYR-- CAENNNC    EINSTSKITCRACRFRKCINAGM
Rm2DBD-NRA3b     CKICGDL-ASGWHCGAITC EACKK FFLRSISTCNG-----YK CQRDLKC LVTKRSRTQCQYCRFQKCISVGM CLICNASASGIHFGAVTC EACKG FFRRS-IK--ENAP-ERYH-- CTENNNC    EIISTSKITCRACRFRKCIEAGM
Rs2DBD-NRA3b     CKICGDI-ASGWHCGAITC EACKK FFLRSISTCNG-----YK CQRDLNC LVTKRSRTQCQYCRFQKCISVGM CLICNASASGIHFGAVTC EACKG FFRRS-IK--ENAP-ERYH-- CAENNNC    EIISTSKITCRACRFRKCIEAGM
Rs1s2DBD-NRA3b   CGAITC EACKK FFLRSISTGDG-----YK CQRDFRC SITKRSRTQCQYCRFQKCISVGM CLICNASASGIHFGTVTC EACKG FFRRS-IK--ENAP-ERYH-- CVENNNC   EIISTSKITCRACRFRKCIEAGM
```

| | | | | | | | | | | |
|---|---|---|---|---|---|---|---|---|---|---|
| **Rs2s2DBD-NRA3c** | CQVCGDL-ASGWHCGAITC | EACKK | FFLRSISTCVG-----YK | CPRDFRC | SITKRSRTQCQYCRFQKCISAGM | CLICNASASGIHFGTVTC | EACKG | FFRRS-IK--ENAP-ERYH-- | CTENNNC | EIISTSKITCRACRFRKCIEAGM |
| **Rso2DBD-NRA3b** | CQICGDL-ASGWHCGAITC | EACKK | FFLRSISTYDG-----YK | CQKELRC | SITKRSRTQCQYCRFQKCISVGM | CLICKASASGIHFGAVTC | EACKG | FFRRS-IK--ENAP-ERYH-- | CTENNNC | EIISTSKITCRACRFRKCIEAGM |
| **Rs2s2DBD-NRA3b** | CQICGDL-ASGWHCGAITC | EACKK | FFLRSISTCDG-----YI | CQRNSTC | SITKRSRTQCQYCRLQKCISVGM | CLICNASASGIHFGVVTC | EACKG | FFRRS-IK--ENAP-ERYY-- | CVENNNC | EIISTSKITCRACRFRKCIEAGM |
| **Rso2DBD-NRA3a** | CQICGDL-ASGWHCGAITC | EACKK | FFLRSISTNDE-----YK | CQKNLNC | SITKRSRTQCQYCRLQKCISVGM | CVICNASASGIHFGAVTC | EACKG | FFRRS-IK--ENAL-ERYH-- | CIENNNC | EIISTSKITCRACRFRKCIQTGM |
| **Rs2s2DBD-NRA3a** | CQICGDL-ASGWHCGAITC | EACKK | FFLRSISTSDE-----YK | CQRNLNC | SITKRSRTQCQYCRLQKCISVGM | CLICNASASGIHFGAVTC | EACKG | FFRRS-IK--ENAP-ERYH-- | CTENNNC | EISTASKITCRACRFRKCIEAGM |
| **Rs1s2DBD-NRA3a** | CQICGDL-ASGWHCGAITC | EACKK | FFLRSISTCDG-----YK | CQRDSRC | SITKRSRTQCQYCRFQKCITIGM | CVICNASASGLHFGAVTC | EACKG | FFRRS-IK--ENAT-ERYH-- | CAENNNC | EIVSTSKIICRACRFRKCIEAGM |
| **Rm2DBD-NRA3a** | CQVCGDL-ASGWHYGAITC | EACKK | FFLRSISPCNG-----LK | CQRNLSC | SMTKRSRTQCQYCRLQKCISVGM | CVICNASASGIHFGAVTC | EACKG | FFRRS-IK--ENAP-DRYY-- | CTENNNC | EIVSTSKTTCRACRFRKCIEAGM |
| **Rs2DBD-NRA3a** | CQVCGDL-ASGWHYGAITC | EACKK | FFLRSISPCNG-----LK | CQRNLSC | SMTKRSRTQCQYCRLQKCISVGM | CVICNASASGIHFGAVTC | EACKG | FFRRS-IK--ENAP-DRYY-- | CTENNNC | EIVSTSKTTCRACRFRKCIEAGM |
| **Dc2DBD-NRA3b** | CQICTDK-ASGWHCGAITC | EACKK | FFLRTINAEN------YK | CQHDKCC | IINRHSRTQCQYCRYQKCLKFGM | CIICSSSSSGLHFGAITC | EACKC | FFRRS-IK--ENAI-EHYH-- | CSANDCC | KIDSKLKMNCRACRFRKCIDGGM |
| **Ar2DBD-NRA1** | CQVCGEQ-SSGLHCGAITC | EACKK | FFLRSINGEDQ-----YK | CVKNKDC | MITRNTRTQCQYCRFQKCLMIGM | CYVCQGPSSGIHFGAITC | EGCKG | FFRRS-IK--ERAP-SRYK-- | CMDNGTC | EMSVSTRNACRYCRFQRCIKVGM |
| **As2DBD-NRA1a** | CQVCGEQ-SSGLHCGAITC | EACKK | FFLRSINGEDQ-----YK | CVKNKDC | MITRNTRTQCQYCRFQKCRIIGM | CYVCQAPSSGIHFGAITC | EGCKG | FFRRS-IK--ERAP-SRYK-- | CMDNGTC | EMTVATRNACRYCRFQRCIKVGM |
| **Rso2DBD-NRA1** | CQVCGEQ-SSGLHCGAITC | EACKK | FFLRSINGEDQ-----YK | CVRNKDC | IITRNNRTQCQYCRFQKCKIIGM | CYVCQAPSSGIHFGAITC | EGCKG | FFRRS-IK--ERAP-SRYK-- | CMDNGTC | EINVATRNACRYCRFQRCIKVGM |
| **Rs2DBD-NRA1b** | CQVCGEQ-SSGLHCGAITC | EACKK | FFLRSINGEDQ-----YK | CVRNKDC | IITRNNRTQCQYCRFQKCKIIGM | CYVCQAPSSGIHFGAITC | EGCKG | FFRRS-IK--ERAP-SRYK-- | CMDNGTC | EINVATRNACRYCRFQRCIKVGM |
| **Rs1s2DBD-NRA1a** | CQVCGEQ-SSGLHCGAITC | EACKK | FFLRSINGEDQ-----YK | CVRNKDC | IITRNNRTQCQYCRFQKCKTIGM | CYVCQAPSSGIHFGAITC | EGCKG | FFRRS-IK--ERAP-SRYK-- | CMDNGTC | EINVATRNACRYCRFQRCIKVGM |
| **Rm2DBD-NRA1** | CQVCGEQ-SSGLHCGAITC | EACKK | FFLRSINGEDQ-----YK | CVRNKDC | IITRNTRTQCQYCRFQKCKIIGM | CYVCQAPSSGIHFGAITC | EGCKG | FFRRS-IK--ERAP-SRYK-- | CMDNGTC | EINVATRNACRYCRFQRCIKVGM |
| **Rs2DBD-NRA1** | CQVCGEQ-SSGLHCGAITC | EACKK | FFLRSINGEDQ-----YK | CVRNKDC | IITRNTRTQCQYCRFQKCKIIGM | CYVCQAPSSGIHFGAITC | EGCKG | FFRRS-IK--ERAP-SRYK-- | CMDNGTC | EINVTTRNACRYCRFQRCIKVGM |
| **Ar2DBD-NRA1b** | CQVCGEQ-SSGLHCGAITC | EACKK | FFLRSINGEDL-----YK | CVRSKDC | VITRNTRTQCQYCRFQKCKFVGM | CYVCQASSSGIHFGAITC | EGCKG | FFRRS-IK--ERAP-SRYK-- | CMDNGTC | EISASTRNGCRYCRFQRCIKVGM |
| **As2DBD-NRA1b** | CQVCGEQ-SSGLHCAAITC | EACKK | FFLRSINGEDL-----YK | CVRNKEC | MITRNTRTQCQYCRFQKCKLIGM | CYVCQAPSSGIHFGAITC | EGCKG | FFRRS-IK--ERAP-SRYK-- | CMDNGTC | EINASTRNGCRYCRFQRCIKVGM |
| **Bc2DBD-NRA1a** | CKVCGEP-SSGWHCGAVTC | EACKK | FFLRSINGEDA-----YK | CIRNQDC | VIVRTTRTQCQFCRFQKCKEVGM | CAVCKAPSSGIHFGAITC | EGCKG | FFRRS-IK--ERAP-ERYR-- | CMENGNC | EISAATRNMCRFCRFQKCLKAKM |
| **Bp2DBD-NRA1** | CKVCGEP-SSGWHCGAVTC | EACKK | FFLRSINGEDA-----YK | CIRNQDC | VIVRTTRTQCQFCRFQKCKEVGM | CAVCKAPSSGIHFGAITC | EGCKG | FFRRS-IK--ERAP-ERYR-- | CMENGTC | EISAATRNMCRYCRFQKCLKAKM |
| **Bc2DBD-NRA1b** | CKVCGEP-SSGWHCGAVTC | EACKK | FFLRSINGEDA-----YK | CIRNKDC | VIVRTTRTQCQFCRFQKCKEVGM | CYVCKSPSSGIHFGAITC | EGCKG | FFRRS-VK--ERAP-ERYR-- | CLENGNC | EISASTRNMCRFCRFQKCISVGM |
| **Bc2DBD-NRA4a** | CRVCGETNSAGWHCGTITC | EACKK | FFLRNVKGDYL-----LK | CIRNSSC | VITKSTRTVCGYCRFQKCFQVGM | CSVCGDSSSGLHFGVITC | EGCKG | FFRRN-IK--L--G-HTFS-- | CSSNGDC | EIGYKTRNACRSCRYKKCISAGM |
| **Bp2DBD-NRA4** | CRVCGETNSAGWHCGTITC | EACKK | FFLRNAKSEYL-----FK | CVRNSSC | VITKSTRTNCAFCRFKKCLQVGM | CSVCGDASSGLHFGAITC | EGCKG | FFRRN-VK--E--G-HKFV-- | CTGDGNC | EISYKSRNACRSCRYNKCVTAGM |
| **Bc2DBD-NRA4b** | CRICGETGTTGWYCGTITC | EACKK | FFMRSVKSDYL-----LK | CVRNSNC | VITKSTRTNCGHCRFQKCLQVGM | CSVCGDASSGFHFGAFTC | EGCKG | FFLRYKNK--K--I-TNES-- | CPNRNIC | QINFSSRNQCKSCRFHKCLTVGM |
| **Bc2DBD-NRA5** | CQVCGDD-NCNWIYGAMIC | EACKK | FFIRSIKEEKR-----YV | CIANKKC | SITKSTRANCQYCRYKKCLDVGM | CVICEQKPSGIHFGVMSC | EACKG | FFRRSCLN--NAS--KQYK-- | CKFDGNGNC | Q-STSRSCRFCRLKKCIEVGM |
| **Bc2DBD-NRA5** | CKVCGDE-ACNWYYGAMVC | EACKK | FFIRSKKEQKR-----YI | CVANKQC | NITKSTRANCQYCRFKKCVQVGM | CAVCSQQPSGIHFGVMSC | EACKG | FFRRSALV--HSD--NPLK-- | CKSNGNQNC | L-LTNRSCRYCRFKKCLEAGM |
| **Ap2DBD-NRB** | CAVCGDQ-ATGRYFGAQIC | **EACKS** | FFIRSTKKGMP-----FK | CQSSGTC | DVTPTSRLLCQYCRFQKCLVAGM | CKVCGDTSSGIHFGVFTC | EGCKG | FFRRS-LK--DG---ASYV-- | CGDNKKC | IITPTSRNVCRYCRYQKCLQVGM |
| **Pmi2DBD-NRB** | CAVCEDQ-ATGRYFGAQIC | EACKS | FFIRSTKKGMP-----FK | CQSSSAC | PVTPTSRLLCQHCRFQKCLNAGM | CKVCNDMSSGIHFGVFTC | EGCKG | FFRRS-LK--DG---ASYM-- | CGDEKKC | VITPTSRNVCRYCRYQKCLQVGM |
| **Aru2DBD-NRB** | CAVCGDQ-ATGRYFGAQIC | EACKS | FFIRSTKKGTP-----FK | CQSSGHC | KVTPTSRLLCQYCRFQKCLTAGM | CQVCGDTSSGIHFGVFTC | EGCKG | FFRRS-LK--DG---TSYF-- | CIDDKKC | LITPTSRNMCRFCRYQKCLQVGM |
| **Lv2DBD-NRB** | CAVCGDQ-ATGRYFGAQIC | EACKS | FFIRSTRKGEP-----FR | CVNNQDC | HITPFSRLLCQYCRYQKCMKAGM | CKVCGDVSSGIHFGVYTC | EGCKG | FFRRS-LR--DR---NTYI-- | CSGKEEC | IITSVTRNHCRYCRFKKCLSVGM |
| **Spur2DBD-NRB** | CAVCGDQ-ATGRYFGAQIC | EACKS | FFIRSTRKGEP-----FR | CVNNQSC | PITPYSRLLCQFCRYQNCMKAGM | CKVCGDVSSGIHFGVYTC | EGCKG | FFRRS-LR--DR---NTYS-- | CSGKEEC | IITPVTRNHCRYCRFKKCLRVGM |
| **Aja2DBD-NRB** | CVVCGDQ-AAGKYFGALIC | EACKS | FFIRSTKKGEP-----FK | CSNNGLC | TITPTSRLLCQCCRYQKCLEVGM | CKVCNDVSSGIHFGVYTC | EGCKG | FYRRS-LR--DS---SNYT-- | CVDNKQC | NITPSTRNVCRYCRFQKCLEVGM |
| **Cg2DBD-NRB1** | CVVCGER-ASGYYFGALVC | **LPCKS** | FYIRCTKDGEPT----FT | CQCNGNC | DIAKQGRIRCQYCRYQRCLMAGM | CKVCGDIANGIHFGVNTC | EGCKK | FFRRG-LV--EN---QSYL-- | CKSEKKC | TINPRNRNNCRYCRYQKCISVGM |
| **Cv2DBD-NRB1** | CVVCGER-ASGYYFGALVC | LPCKS | FYIRCTKDGEPT----FT | CQCNGNC | DIAKQGRIRCQYCRYQRCLMAGM | CKVCGDIANGIHFGVNTC | EGCKK | FFRRG-LV--EN---QSYL-- | CKSEKKC | TINPRNRNNCRYCRYQKCISVGM |
| **My2DBD-NRB1** | CVVCGEK-ASGYYFGALVC | LPCKS | FYIRCTKEGEPT----FT | CQCNGNC | DIAKQGRIRCQYCRYQRCLMAGM | CKVCGDIANGIHFGVNTC | EGCKK | FFRRG-LV--EN---QSYV-- | CKGDKKC | TINPRNRNNCRFCRYQKCITVGM |
| **Pm2DBD-NRB1** | CVVCGEK-ASGYYFGALVC | LPCKS | FYIRCTKEGEPT----FT | CQCNGNC | DIAKQGRIRCQYCRYQRCLMAGM | CKVCGDIANGIHFGVNTC | EGCKK | FFRRG-LV--EN---QSYV-- | CKGDKKC | TINPRNRNNCRFCRYQKCITVGM |
| **Mc2DBD-NRB1** | CVVCGEK-ASGFYFGALVC | LPCKS | FYIRCTKDGEPT----FT | CQCNGNC | DIAKQGRIRCQYCRYQRCLMAGM | CKVCGDIANGIHFGVNTC | EGCKK | FFRRG-LV--EN---QGYN-- | CKGEKSC | QINPRNRNNCRYCRYQKCISAGM |
| **MgMR7B1** | CVVCGEK-ASGFYFGALVC | LPCKS | FYIRCTKDGEPT----FT | CQCNGNC | DIAKQGRIRCQYCRYQRCLMAGM | CKVCGDIANGIHFGVNTC | EGCKK | FFRRG-LV--EN---QGYN-- | CKGEKSC | QINPRNRNNCRYCRYQKCISAGM |
| **Me2DBD-NRB1** | CVVCGEK-ASGFYFGALVC | LPCKS | FYIRCTKDGEPT----FT | CQCNGNC | DIAKQGRIRCQYCRYQRCLMAGM | CKVCGDIANGIHFGVNTC | EGCKK | FFRRG-LV--EN---QGYN-- | CKGEKSC | QINPRNRNNCRYCRYQKCISAGM |
| **Mm2DBD-NRB1** | CVVCGER-ASGFYFGALVC | LPCKS | FYIRCTKDGEPT----FT | CQCSGNC | DIVKQGRIRCQYCRYQRCLMAGM | CKVCGDIANGIHFGVNTC | EGCKK | FFRRG-LV--EN---QSYI-- | CKGEKEC | GINPRNRNNCRYCRYQKCLTVGM |
| **Po2DBD-NRB3** | CDVCGEQ-ASGHYFGALVC | LPCKS | FFIRCTKDGRPT--FSTQ | C--GGKC | DILKGGRVRCQHCRFQKCINAGM | CKVCGDIANGIHFGVTTC | EGCKK | FFRRG-LK--EN---RSYT-- | CKGSMHC | SINPRMRNNCRYCRYQKCLLEGM |
| **Ema2DBD-NRB3** | CDVCGEQ-ASGHYFGALVC | LPCKS | FFIRCTKDGRPS--FSSQ | C--GGKC | DVLKGGRVRCQHCRFQRCISAGM | CKVCGDIANGIHFGVTTC | EGCKK | FFRRG-LK--EN---RSYT-- | CKGSMHC | SINPRMRNNCRYCRYQKCLLEGM |
| **Ech2DBD-NRB3** | CDVCGEQ-ASGHYFGALVC | LPCKS | FFIRCTKDGRPS--FSSQ | C--GGKC | DVLKGGRVRCQHCRFQRCISAGM | CKVCGDIANGIHFGVTTC | EGCKK | FFRRG-LK--EN---RSYT-- | CKGSMHC | SINPRMRNNCRYCRYQKCLLEGM |
| **Bg2DBD-NRB3** | CDVCGEQ-ASGHYFGALVC | LPCKS | FFIRCTKDGRPN--FSNQ | C--GGKC | DVLKGGRVRCQHCRFQKCIHAGM | CKVCGDIANGVHFGVTTC | EGCKK | FFRRG-LK--EN---KSYT-- | CKANMHC | SINPRMRNNCRYCRYQKCLYEGM |
| **Bt2DBD-NRB3** | CDVCGEQ-ASGHYFGALVC | LPCKS | FFIRCTKDGRPN--FSNQ | C--GGKC | DVLKGGRVRCQHCRFQKCISAGM | CKVCGDIANGVHFGVTTC | EGCKK | FFRRG-LK--EN---RTYT-- | CKASQMC | SINPRMRNNCRYCRYKKCITEGM |
| **Aca2DBD-NRB3** | CDVCGEQ-ASGHYFGALVC | LPCKS | FFIRCTKDGHPN--FSNQ | C--GGKC | DVLRGGRVKCQHCRFQKCISAGM | CKVCGDIANGVHFGVTTC | EGCKK | FFRRG-LK--EN---RSYT-- | CKGSMRC | SINPRMRNNCRYCRYQKCLFEGM |
| **Pca2DBD-NRB3** | CDVCGEP-ASGHYFGALVC | LPCKS | FFIRCTKTGDPV--FSSQ | C--GGTC | DVHKLGRIRCQFCRFQRCLSAGM | CKVCCDIANGVHFGVTTC | EGCKK | FFRRG-LK--EH---QAYV-- | CKVAKKC | TINPRMRNNCRYCRFQKCLNVGM |
| **Pvd2DBD-NRB3** | CDVCGEP-ASGHYFGALVC | LPCKS | FFIRCTKTGDPV--FSSQ | C--GGTC | DVHKLGRIRCQFCRFQRCLSAGM | CKVCCDIANGVHFGVTTC | EGCKK | FFRRG-LK--EH---QAYV-- | CKVAKKC | TINPRMRNNCRYCRFQKCLNVGM |
| **Po2DBD-NRB2** | CVVCHEA-ASGNFFGAVVC | LPCKS | FFIRCTKDSESC--IVRQ | C--RGQC | DISKQLRNRCQFCRYQRCLAAGM | CRVCGDLSNGIHFGVFTC | EGCKK | FFRRG-LK--EH---TTYV-- | CKEHGMC | RLTPRNRNSCRRCRYRKCIDVGM |
| **Aca2DBD-NRB2** | CVVCGEV-ANGNFFGAVVC | LPCKS | FFIRCTKDGEPC--ILQQ | C--GGNC | DIKKQLRNRCQYCRYQQCLAMGM | CKVCGDLANGIHFGVYTC | EGCKK | FFRRG-LK--EH---LSYV-- | CKDAKRC | RLNPRNRNNCRYCRYQKCLLVGM |
| **Hruf2DBD-NRB2** | CQVCGEK-ASGNFFGALVC | LPCKS | FFIRCTKEGEPV--FVSQ | C--DSRC | DVSKQGRNRCQHCRYKKCLAAGM | CRVCGDIANGIHFGVYTC | EGCKK | FFRRG-LL--EY---QSYV-- | CKGAMSC | LLNPRNRNNCRYCRYQKCLNVGM |
| **Hrub2DBD-NRB2** | CQVCGEK-ASGNFFGALVC | LPCKS | FFIRCTKDGEPV--FVSQ | C--DGRC | DVSKQGRNRCQNCRYKKCLAAGM | CRVCGDIANGIHFGVYTC | EGCKK | FFRRG-LL--EY---QSYV-- | CKGAMLC | VLNPRNRNNCRYCRYQKCLNVGM |
| **Pca2DBD-NRB2** | CAVCGDR-ASGNFFGATVC | LPCKS | FFIRCTKDGEPA--VQKQ | C--NGCC | DVSKELRNRCQFCRFQRCLLIGM | CRVCGDLANGVHFGIFTC | EGCKK | FFRRG-LK--EH---KSYV-- | CRGAHDC | QLNPRNRNNCRACRFWKCAAVGM |
| **Ga2DBD-NRB1** | CAVCGEE-ANGNFFGALVC | LPCKS | FFIRCNKANELTS-VQRP | C--DNRC | TATRQGRVRCQRCRYLKCVQVGM | CLVCGDIANGIHFGIKTC | EGCKK | FFRRG-LL--EN---KTYV-- | CTKQQAC | IINPRNRNNCRLCRYRKCVSSGM |
| **Lg2DBD-NRB1** | CVVCKEP-AHGYNFGALVC | LPCKS | FYIRCSKEDVIY----SK | C--RGQC | DITGENRIRCQYCRYQQCLKVGM | CMVCGDLSNGIHFGVGTC | EGCKK | FFRRC-LL--ES---SKLI-- | CVNERRC | KINPKTRNRCRLCRYLKCLQVGM |
| **Hruf2DBD-NRB1** | CVVCADN-ASGQYFGATVC | LPCKS | FYIRCTKEGEPV--FSSK | C--HGSC | DVTKQARIRCQSCRYQKCLKAGM | CLVCGDLANGIHFGVYTC | EGCKK | FFRRG-LV--EN---RTYQ-- | CKGDNTC | VINPRSRNTCRACRYRKCLSVGM |
| **Pau2DBD-NRB** | CSVCGAD-ASGMYFGAIVC | LPCKS | FFIRCTKEGEPTR---LK | CQNDINC | GYPGPIRHRCQYCRFQRCLVAGM | CKVCGDIANGIHFGVNTC | EGCKK | FFRRG-LK--EA---DTYA-- | CKEGQAC | PITPRSRNACRFCRYMKCVTVGM |
| **Ct2DBD-NRB1** | CVVCEDK-AEGRYFGATVC | LPCKS | FFIRCTKEGPKR----LV | CHATKNC | DMTGGNRMRCSFCRHQKCLSMGM | CRVCSDIANGVHFGVTTC | EGCKK | FFRRA-LK--EY---RTYR-- | CKFNLMC | TVNPRNRNTCRSCRYQKCIRLGM |
| **Ls2DBD-NRB1** | CAVCSEK-ANGRYFGAIVC | LPCKS | FFIRCTKDGKPT----LS | CPENMAC | DITGVNRMRCQYCRYKKCLSVGM | CRVCLDIANGIHFGVTTC | EGCKK | FFRRA-LK--EY---LMYD-- | CKFLKHC | FINPRNRNSCRYCRYKKCLGVGM |
| **La2DBD-NRB** | CTVCDEP-ASGIYFGALVC | **LACKS** | FYIRCTREGHKTD---YR | CAAVGRC | LLDKPYRVRCQQCRFQKCLSVGM | CKVCGYMANGVHFGVETC | EGCKK | FFRRG-LK--EC---ATYH-- | CKALKDC | AINPKNRNDCRYCRYQKCLAVGM |
| **Hr2DBD-NRB2** | CVVCGDS-AHGNYFHAFVC | **VPCKT** | FFLRYADEHAQ-----FQ | CKFNGNC | EITLTTRTNCKHCRYQKCVKQGM | CVVCGDLANGIHFGANTC | EGCKK | FFRRG-LT--ES---GSYS-- | CKNNHDC | KINPNTRNNCRFCRFKKCLDVGM |
| **Ls2DBD-NRB2a** | CKVCSDT-ATGMYFGALVC | VPCKT | FFLRYAGDNFA-----FR | CKRDKGC | EIVTATRTQCKYCRYQKCLHVGM | CVVCLDLANGVHFGAVTC | EGCKK | FFRRG-LT--EH---ESYQ-- | CKLDKSC | PINPRTRNACRYCRYQKCINVGM |
| **Ls2DBD-NRB2b** | CQVCGEK-ASGNFFGALVC | **LPCKT** | FFIRCSSEEVST----FR | CRVDFNC | EVTGSRRSKCKYCRYRKCIAVGM | CVVCHDVANGVHFGIVSC | EGCKK | FFRRG-LQ--ES---ESYI-- | CKQNGAC | VINPKTRNDCRYCRFQKCLMAGM |
| **Ct2DBD-NRB2** | CQVCNDH-AAGTYFGAKVC | **IPCKA** | FFIRSTSDKTSV----FE | CPGDQRC | KITVMTRTACKFCRFQNCLKAGM | CVICGDDANGVHFGALTC | EGCKK | FYRRG-LLVGEA---KSYI-- | CKADKTC | EITARTRNNCRYCRFQKCLRMGM |
| **Of2DBD-NRB2** | CSVCGSR-AAGVFFGVLTC | WTCKT | FFIRHQKAGAGG----LK | CDNNGQC | KLDLKLRRRCAYCRYKKCIEVGM | CLVCGDIASGVHFGIVAC | LGCTK | FFRRT-IRMGTE---DSYM-- | CYNAHRC | VINPQNRNTCPRCRLDKCKRLGM |
| **Cb2DBD-NRC** | CKVCEKP-SNGLHYGVRAC | DSCKC | FFARSLKRSTES----LV | CSKDSNC | NFGRQPKTICSYCRFQKCVKVGM | CKVCEKPGHAIHYGVWTC | DSCKW | FFTNS-LR---RGT-ESLF-- | CHKGSSC | DLSSNPFETCKYCRFQKCVKLGM |
| **Cr2DBD-NRC1** | CKICGGY-ARGVNYGVLSC | DNCRA | FFSHYLHRKK------LK | CRKNNNC | VIDKFTSNKCGKCRMEKCLRMGM | CKVCGDVPVGIGYGVLSC | GSCRM | FFRRH------NG--TDVHLS | CNKEGKC | DL-RLKFAKCPKCRMRKCLEVGM |
| **Cr2DBD-NRC2** | CKICGGY-AKGTNYGVLTC | DGCRT | FFIQYIHMKE------LK | CRRNNNC | VIDKFITKKCAKCKIEKCLIMGM | CKVCGDTASKIFYGVLTC | EGCRV | FFRRY------AG--KDVKLM | CYSRGNC | NL-LEKFTKCQKCRMRKCLEVGM |
| **Cr2DBD-NRC3** | CKVCGDK-PRGVHYGVLAC | EGCRS | FFRTNRDKKE------LK | CRKSNKC | VVDKYSRNKCGKCRMEKCLKLGM | CRVCGDISFEHFFDIVAC | EGCKG | FFRTH------HE--KDIKLE | CRVDGNC | DINIISRNTCKSCRMKKCLEAGM |
| **Cr2DBD-NRC4** | CNICGDV-ANGVHYGVPAC | EGCKK | FFSRMHNKQEY-----FL | CRKGNNC | FIDRESRNSCQSCRLAKCRQAGM | CEVCGDERYSTNRVPAYC | EACKT | VYSSH------IK--EIKMFQ | CEKDGKC | VINKWTRDKCQSCRMGKCLEAGM |
| **Aa2DBD-NRC3** | CVICADL-ANAYHYGVASC | NGCKT | FFRRTIVDGHAD----LR | CQFEGHC | EVSKEIRNACRRCRFDKCLQAGM | CLICSDVATGYHYGVTSC | HGCKA | FFRRTIIA---G---RTF--V | CDRDGTC | AVTKGDPTFCRGCRLTKCLEAGM |
| **Aa2DBD-NRC4** | CVICADL-ANAYHYGVASC | NGCKT | FFRRTIVDGHAG----LR | CQFEGRC | EVSKDGRNACRYCRFNKCLQAGM | CLICSDVATGYHYGVASC | HGCKA | FFRRSIIA---G---RTF--V | CDRGGAC | AVMKGDPTFCRGCRLTKCLEAGM |
| **Aa2DBD-NRC5** | CAICSDL-ANAYHYGVASC | NGCKT | FFRRTIVDGHAG----LR | CQFEGRC | EVSKEVRNACRRCRFDKCVLAGM | CLICSDFATGYHFGASSC | HGCKS | FFRRTIVA---G---RNF--V | CERDGTC | AVQKRDSTSCRGCRFNKCLEAGM |
| **Aa2DBD-NRC2** | CAICNDI-ASGYHFGAASC | AGCRI | FFRRNVLSGDSE----LR | CHYEGNC | EVSKDVRTACRHCRFNKCLLAGM | CAICSDVATGYHYGVAAC | FGCKT | FFRRTVVA---G---HTY--I | CQHGGTC | VVNKDQRMACRHCRLNKCIQAGM |
| **Aa2DBD-NRC1** | CAICDDT-ASGTHYGVTSC | TGCKT | FFRRTIVDGHAE----LR | CQYEGHC | VVSKDVRNACRQCRFDKCLLAGM | CAICEDVATGYHYGVAAC | LGCKA | FFRRTIVA---G---HAY--T | CQRDGNC | IVNVDGRMACRHCRFNKCLQAGM |
| **Aa2DBD-NRC6** | CAICDGT-ASGYHYDVASC | TACWM | FFRQAVVSGEAE----LR | CEYQGNC | IVSKDVRNACRHCRFNKCLLEGM | CAICDDVANGYHYGVASC | LGCKG | FFRRTIVS---G---DAARIR | CQHDGNC | DVNKRNRNACRRCRFDKCQLAGM |
| **Aa2DBD-NRC7** | CVICDAT-ASGYHYDVASC | TACWM | FFRQMVITGEAE----LR | CQRQGNC | VVSKDVRNACRHCRFNKCLLEGM | CAICHDAATGYHYGVAAC | HACKT | FFRRVIVM---G---DSARIR | CQKDGNC | DVSKRNRNACRRCRFDKCQQAGM |
