## Supplemental material 3 for "Updated knowledge and a proposed nomenclature for nuclear receptors with two DNA Binding Domains (2DBD-NRs)"

Supplemental material 3. Ti sequence for sequence logo

(NCBI accession number of 2DBD-NRs see table 6, accession number of other NR is in the bracket after each NR name)

```
2DBD-NRA
Bf2DBD-NRA                    FAK-KIPGFRTCSID-DQISMIQ
Bf2DBD-NRA                    FAK-KIPGFRTCSID-DQISMIQ
OfuNR7A1                      YAK-KIPGFREFPVE-DQILLLQ
Sb2DBD-NRA2                   FAK-LVPGFNQLGIT-ARSNLVR
Sm2DBD-NRA2                   FAK-LVPGFNQLGIT-ARSNLVR
Sj2DBD-NRA2                   FAK-LVPGFNQLGIT-ARSNLVR
Of2 BD-NRA2                   FAK-LVPGFNQLGIT-ARSNLVR
Ov2DBD-NRA2                   FAK-LVPGFNQLGIT-ARSNLVR
Cs2DBD-NRA2                   FAK-LVPGFNQLGIT-ARSNLVR
Pw2DBD-NRA2                   FAK-LVPGFNHLGIA-ARSNLVR
Pv2DBD-NRA2                   FAK-LVPGFNHLGIA-ARSNLVR
Ph2DBD-NRA2                   FAK-LVPGFNHLGIA-ARSNLVR
Psm2DBD-NRA2                  FAK-LVPGFNHLGIA-ARSNLVR
Fh22DBD-NRA2                  FAK-LVPGFNHLGIT-ARSNLVR
Eg2DBD-NRA2                   FAK-LIPGFNQMSLS-ARGHLVR
Em2DBD-NRA2                   FAK-LIPGFNQMSLS-ARGHLVR
Hm2DBD-NRA2                   FAK-LIPGFNQMSLS-ARGHLVR
Eg2DBD-NRA2                   FAK-LIPAFNQMSLS-ARGHLVR
Ta2DBD-NRA2                   FAK-LIPGFNQMSLS-ARGHLVR
Hd2DBD-NRA2                   FAK-LIPGFNQMSLS-ARGHLVR
Mc2DBD-NRA2                   FAK-LIPGFNQLSLA-GRGNLVR
Se2DBD-NRA2                   FAK-LIPGFNQLELT-ARGALVR
Sme2DBD-NRA3                  FVK-AIPGFNKFELK-SRRSLVQ
Sj2DBD-NRA3                   FVK-YIPGFCYLKIS-DQRQLVR
Sm2DBD-NRA3                   FVK-YIPGFCYLKIS-DQRQLVR
Ph2DBD-NRA3                   FVK-LIPGFNQLVLD-DRRQLVR
Pw2DBD-NRA3                   FVK-LIPGFNQLVLD-DRRQLVR
Of2DBD-NRA3                   FVK-LIPGFDQLELG-DKRQLVR
Ov2DBD-NRA3                   FVK-LIPGFDQLELG-DKRQLIR
Em2DBD-NRA3                   FSK-LIAGFNRLGIN-DRRQLIR
Eg2DBD-NRA3                   FSK-LIAGFNRLGIN-DRRQLIR
Ht2DBD-NRA3                   FSK-LIAGFNRLGIN-DRRQLIR
Ts2DBD-NRA3                   FSK-LIAGFNRLGIN-DRRQLIR
Hs2DBD-NRA3                   FSK-LIAGFNGLGIN-DRRQLIR
Hm2DBD-NRA3                   FSK-LIAGFNGLGIN-DRRQLIR
Spr2DBD-NRA3                  FTK-IIPGINRLELN-DKRQLVR
Sme2DBD-NRA3                  FAK-AVPGFKELFHN-DMKVLVQ
Sj2DBD-NRA1                   FAQ-TISSCQELSEF-DMKILIQ
Sj2DBD-NRA1                   FAQ-TISSCQELSEF-DMKILIQ
Sma2DBD-NRA1                  FAQ-SIPSFQELSQY-DMKILIQ
Sh2DBD-NRA1                   FAQ-SIPSFQDLSEY-DMKILIQ
Sm2DBD-NRA1                   FAQ-SIPNFQELSEF-DMKILIQ
Fb2DBD-NRA1                   FAR-AVPGFRDLSRM-DMKTLVQ
Fh2DBD-NRA1                   FAR-AVPGFRDLSRL-DMKTLVQ
Pw2DBD-NRA1                   FAR-AVPGFRDLSRL-DMKTLVQ
Pw2DBD-NRA1                   FAR-AVPGFRDLSRL-DMKTLVQ
Cs2DBD-NRA1                   FAR-AVPGFRDLPSV-VMKKLVQ
Of2DBD-NRA1                   FAR-AVPGFRELPSV-VMKKLVQ
Em2DBD-NRA1                   FAR-AIPGFCDLPRA-DTKFLLQ
Eg2DBD-NRA1                   FAR-AIPGFCDLPRA-DTKFLLQ
Hm2DBD-NRA1                   FAR-AIPGFRDLPRA-DTKFLVQ
Hn2DBD-NRA1                   FAR-AIPGFRDLPRA-DTKFLVQ
Hd2DBD-NRA1                   FAR-AIPGFRDLPRA-DTKFLVQ
Mc2DBD-NRA1                   FAR-AIPGFRDLPRA-DTKFLVQ
Ts2DBD-NRA1                   FAR-AIPGFRDLPRA-DTKFLVQ
Sp2DBD-NRA1                   FAR-AIPGFRDLPRS-DTKVLVQ
Cv2DBD-NRA                    ALK-MFPVFKILELD-DRITLVQ
Cg2DBD-NRA                    ALK-MFPVFKILELD-DRITLVQ
Pm2DBD-NRA                    FSK-KVPGFRALSLD-DQIKLVQ
Mc2DBD-NRA                    FAK-KCQPFRKLALE-DQVRMLQ
Mg2DBD-NRA                    FAK-KCQPFRKLALE-DQVRMLQ
Dp2DBD-NRA                    FAK-LVPGFKGLCLN-DQVKLIQ
Me2DBD-NRA                    FAK-KCQPFRKLALE-DQVRMLQ
Mm2DBD-NRA                    FAK-LVPGFKTLSLN-DQVKLIQ
Po2DBD-NRA                    YAK-KVPGFRDIKLD-DQVRLLS
Em2DBD-NRA                    FAK-KVPGFRDLKLD-DQVKLLR
Hruf2DBD-NRA                  FSK-KLPNFQSLSLQ-DQVSLIQ
My2DBD-NRA                    FSK-KVPGFRAISLD-DQIKLIQ
Pca2DBD-NRA                   FAK-RVPGFRKLPLD-DQVLLVQ
Ech2DBD-NRA                   YAK-KTSGFRDLKLD-DQVKLLS
Bg2DBD-NRA                    FAK-KVPGFRLLKIE-DQVLLLR
Ga2DBD-NRA                    FAK-KVPGFKSLVIE-DQILLMQ
Hrub2DBD-NRA                  FSK-KLPNFQSLLLE-DQVSLIQ
Acas2DBD-NRA                  FAK-KICGFRSLDIN-DQVLLLR
Bt2DBD-NRA                    FAK-KVPGFRQLKIE-DQVLLLR
Bat2DBD-NRA                   FAK-KVPDFRKLPID-DQVALVQ
Ng2DBD-NRA                    FAK-KIPGFR-KLSISDQITLVQ
La2DBD-NRA2                   FGK-KIPGFR-AIPMEDQICLVR
Bc2DBD-NRA4a                  FIS-NIPYITKFSDI-DKNIIYV
Bc2BD-NRA4b                   FIK-QTKFLRKFNEN-DQNIILI
Bp2BD-NRA4                    -QE-HELVLERFSIE-DQKILLV
Ar2DBD-NRA2a                  FVD-GIPNFNSLNVE-DKTSLVY
Ar2DBD-NRA2b                  FIE-GIPNFNSLNVE-DKTSLVY
Rs2s2DBD-NRA2b                FID-RIPNFNSIHFI-EKTSLTH
```

| | |
|---|---|
| Rso2DBD-NRA2b | FID-KIPNFNSISFI-EKTSLIH |
| Rs1s2DBD-NRA1a | FID-KIPNFNSINFI-EKSSLIH |
| Rm2DBD-NRA2b | FID-RIPNFNSINFI-EKTSIIH |
| Rso2DBD-NRA2a | FAL-RIPDFCSLNSE-DKNLLIH |
| Rs1s2DBD-NRA2d | FAL-RIPDFCTLNSE-DKSLLIH |
| Rm2DBD-NRA2a | FAL-RIPEFCSLTAD-DKNLLIH |
| Ar2DBD-NRA2a | FAL-TIPEFSSLNSD-DRTLLIH |
| As2DBD-NRA2 | FTL-NITDFCILNSD-DKNLLID |
| Dc2DBD-NRA2b | FST-EIPGFSTFAHS-DHLLLIR |
| Dc2DBD-NRA2a | FAT-EIPGFSTFIHD-DNLLLIR |
| Rso2DBD-NRA3a | FCQRTIPGFTNMS---NKYEIIS |
| Rs2s2DBD-NRA3a | FCQSTIPDFTNMP---DKYEIIS |
| Rsp12DBD3 | FCQKTIPGFTNMS---QKYEIIS |
| Rsp2_2D6 | FCQRTIPGFINMS---QKYEIIS |
| Rm2DBD-NRA3b | FCQSIIPGFSNTS---DKYEIIS |
| Rm2DBD-NRA3a | FCQKIIPDFTSMP---DQWEIIS |
| Ar2DBD-NRA3a | FCQKTIPGFDSMY---EKYDVIS |
| As2DBD-NRA3a | FCQKTIPGFDNMY---EKYEVIS |
| Ar2DBD-NRA3b | FCQKTIPGFDNMY---EKYDVVS |
| As2DBD-NRA3b | FCQKTIPGFDNIY---EKYEIIS |
| Dc2DBD-NRA3a | FCQKTIYGFSELS---DPYEVLS |
| Dc2DBD-NRA3b | FCQKTITNFNNLS---DPYEIIS |
| Bc2DBD-NRA1b | FLS-DFKEFMNFNQM-DREELIK |
| Ar2DBD-NRA1b | FAT-RIPGFMNLQNVTDQVQLIK |
| As2DBD-NRA1b | FAT-RIPGFMNLQNVTDQVQLVK |
| Rso2DBD-NRA1 | FAT-RIPGFMNFHNVNDQIQLIK |
| Ar2DBD-NRA1a | FAT-RIPNFMNFHTVTDQIQLIK |
| **2DBD-NRB** | |
| Pmi2DBD-NRB | FAK-NIPGFREIPVS-DQMNLIK |
| Aru2DBD-NRB | FAK-NIPGFCHISVS-DQMSLIK |
| Ema2DBD-NRB | FCR-NIPGYFKIDIQ-DRISLGK |
| Ap2DBD-NRB | FAK-NIPGFRHISVN-DQMNLIK |
| Aca2DBD-NRB | FCR-NIPGYFKLDIQ-DRICLGK |
| Lv2DBD-NRB | YIK-KIPGFASIDMN-DRINLIK |
| Cg2DBD-NRB | FVK-TIPGFDSINSE-DAAELCV |
| Spur2DBD-NRB | YIK-KIPGFPSIHMN-DRLYLIK |
| Cv2DBD-NRB | FVR-TIPGFESINQE-DAAELCI |
| Hruf2DBD-NRB2 | FMS-KVPGFKDLNPE-DRHFVVQ |
| Mm2DBD-NRB1 | FAS-QIPGFSNLHPE-DKAFLCK |
| Pca2DBD-NRB3 | CLA-ALPGCEDLTSE-DRALLSQ |
| Aca2DBD-NRB2 | FVL-KLPGMDKIHPK-DRVKLGN |
| Hrub2DBD-NRB2 | FMS-KVPGFKDLNPE-DRNFVVQ |
| Me2DBD-NRB1 | FVM-QIPGFMQLDHD-DKQDLCQ |
| Mc2DBD-NRB1 | FCM-QIPGFMQLDHE-DKQDLCQ |
| Ech2DBD-NRB3 | FCR-NIPGYFKIDIQ-DRISLGK |
| **2DBD-NRC** | |
| Cb2DBD-NRC | FLR-GIPRMDEIEME-ERLMLLK |
| Cr2DBD-NRC2 | FVR-NLPHFNLFGMN-DKIILFK |
| Cr2DBD-NRC1 | FVN-RLPSIDSFEIS-DKAILFK |
| Cr2DBD-NRC3 | FVN-HLPFIKTFEIN-DKVILFK |
| Cr2DBD-NRC4 | FAN-HLPFIKTFEIS-DKTILFK |
| **NRs from Subfamily 1** | |
| hTRa (NP_955366.1) | FAK-KLPMFSELPCE-DQIILLK |
| mTRa (CAA30577.1) | FAK-KLPMFSELPCE-DQIILLK |
| hTRb (NP_000452.2) | FAK-KLPMFCELPCE-DQIILLK |
| mTRb (NP_033406.1) | FAK-KLPMFCELPCE-DQIILLK |
| mRARa (CAA71177.1) | FAK-QLPGFTTLTIA-DQITLLK |
| hRARa (NP_000955.1) | FAK-QLPGFTTLTIA-DQITLLK |
| hRARb (NP_000956.2) | FAK-RLPGFTGLTIA-DQITLLK |
| hRARg (NP_000957.1) | FAK-RLPGFTGLSIA-DQITLLK |
| mRARg (NP_001398643.1) | FAK-RLPGFTGLSIA-DQITLLK |
| hRORa (NP_599023.1) | FAK-RIDGFMELCQN-DQIVLLK |
| hRORb (CAA69929.1) | FAK-RITGFMELCQN-DQILLLK |
| mRORg (NP_035411.2) | FAK-RLSGFMELCQN-DQIILLK |
| hRORg (AAA64751.1) | FAK-RLSGFMELCQN-DQIVLLK |
| dHR3 (NP_001246236.1) | FAK-LIPGFMRLSQD-DQILLLK |
| dECR (NP_001163061.1) | FAK-GLPAFTKIPQE-DQITLLK |
| mLXRb(NP_001272446.1) | FAK-QVPGFLQLGRE-DQIALLK |
| hLXRb (BAH02288.1) | FAK-QVPGFLQLGRE-DQIALLK |
| hLXRa (NP_001171201.1) | FAK-QLPGFLQLSRE-DQIALLK |
| hFXRa (NP_005114.1) | FTK-KLPGFQTLDHE-DQIALLK |
| mFXRa (NP_033134.2) | FTK-KLPGFQTLDHE-DQIALLK |
| hVDR (XP_024304946.1) | FAK-MIPGFRDLTSE-DQIVLLK |
| hPXR (BAH02292.1) | FAK-VISYFRDLPIE-DQISLLK |
| mPXR (NP_035066.1) | FAK-VISYFRDLPIE-DQISLLK |
| hCAR (NP_001070948.1) | FTK-DLPVFRSLPIE-DQISLLK |
| mCAR (AAC53349.1) | FTK-DLPLFRSLTME-DQISLLK |
