## Supplemental material 4 for "Updated knowledge and a proposed nomenclature for nuclear receptors with two DNA Binding Domains (2DBD-NRs)"

Supplemental material 4. AF2 sequence (yellow highlighted) for sequence logo

(NCBI accession number of 2DBD-NRs see table 6, accession number of other NR is in the bracket after each NR name)

```
2DBD-NRA
Rso2DBD-NRA1                   IEFPAFFSRVYLNEQDI
Ar2DBD-NRA1a                   VELPAFFLRVYLNEQD-
Ar2DBD-NRA1b                   ITYPAFFSRVYLNEHDT
As2DBD-NRA1b                   IEYPAFFSRVYLNEHNI
Rso2DBD-NRA3a                  RNLPSFFYRIFLP----
Rsp2DBD-NRA3a                  GNLPSFFYRIFLP----
Rs2s2DBD-NRA3a                 PNLPSFFYSIFLP----
Rm2DBD-NRA3a                   QNLPSFFYSIFLP----
Rs1s2DBD-NRA3b                 RNLPSFFYRIFVPKQIG
Rs2s2DBD-NRA3c                 RNLPSFFYRIFVP----
Rm2DBD-NRA3b                   RNLPSFFYRIYVP----
Rso2DBD-NRA3b                  RNLPSFFYRIYVP----
Ar2DBD-NRA3b                   QNVPSYFYRIYLP----
Ar2DBD-NRA3a                   RNLPSFFYRIYVP----
As2DBD-NRA3a                   RNLPSFFYRIYVP----
As2DBD-NRA3b                   PNLPSFFYPIYLP----
Dc2DBD-NRA3a                   RNLPSFFYRIYVPN---
Dc2DBD-NRA3b                   KNLPFYFYRIYVP----
Sb2DBD-NRA2                    LRFPELYVEMFQLADSA
Sm2DBD-NRA2                    LRFPELYVEMFQLADSA
Sj2DBD-NRA2                    LRFPELYVEMFQLADSA
Of2DBD-NRA2                    LHFPELYIEMFQLADSA
Cs2DBD-NRA2                    LHFPELYIEMFQLADSA
Ov2DBD-NRA2                    LHFPELYIEMFQLADSA
Pw2DBD-NRA2                    LHFPELYVEMFQLADSA
Phe2DBD-NRA2                   LHFPELYVEMFQLADSA
Psm2DBD-NRA2                   LHFPELYVEMFQLADSA
Fh2DBD-NRA2                    LYFPELYVEMFRLADDA
Eg2DBD-NRA2                    LRYPDLYVEMYEIGESA
Eg2DBD-NRA2                    LRYPDLYVEMYEIGESA
Em2DBD-NRA2                    LRYPDLYVEMYEIGESA
Ta2DBD-NRA2                    LRYPDLYVEMYEIGESA
Hm2DBD-NRA2                    LRYPDLYSEMYELPR
Hd2DBD-NRA2                    LRYPDLYAEMYEINETS
Mc2DBD-NRA2                    LGYPDLYVEMYQLTDPP
Se2DBD-NRA2                    LHFPELFVEMFQLSVTP
Sj2DBD-NRA2                    LEFPQLYLEMFQLTDEE
Sm2DBD-NRA3                    LEFPQLYLEMFQLTDEE
Ph2DBD-NRA3                    LVFPELYTEMFQLTEEA
Pw2DBD-NRA3                    LVFPELYTEMFQLTEEA
Of2DBD-NRA3                    LDFPELYTEMFQLTEAE
Ov2DBD-NRA3                    LDFPELYTEMFQLTEAE
Em2DBD-NRA3                    LVFPELYVEMFAIGADK
Eg2DBD-NRA3                    LVFPELYVEMFAIGADK
Ts2DBD-NRA3                    LVFPELYVEMFAIGADK
Hs2DBD-NRA3                    LIFPELYLEMFAIGAGE
Hm2DBD-NRA3                    LIFPELYLEMFAIGAGE
Ht2DBD-NRA3                    LVFPELYVEMFAIGTDK
Spr2DBD-NRA3                   LVFPALFTEMFALDEPI
Sj2DBD-NRA3                    QVFPDFYMQLFQLDD--
Sma2DBD-NRA3                   LPFSDLYIQLFQLNE--
Sh2DBD-NRA3                    LPFSDLYIQLFQLNE--
Sm2DBD-NRA1                    LPFSNLYIQLFQLNE--
Fb2DBD-NRA1                    LTFPDLYVQMFRLDE--
Fh2DBD-NRA1                    LTFPDLYVQMFRLDE--
Pw2DBD-NRA1                    LVFPDLYVQMFQLND--
Pw2DBD-NRA1                    LVFPDLYVQMFQLND--
Cs2DBD-NRA1                    LKFPDLYVQMFEL-D--
Of2DBD-NRA1                    LKFPDLYVQMFEL-D--
Em2DBD-NRA1                    LHFPDLFVQMFQLED--
Eg2DBD-NRA1                    LHFPDLFVQMFQLED--
Ts2DBD-NRA1                    LHFPDLFVQMFQLGS--
Mc2DBD-NRA1                    LHFPDLFVQMFQLDV--
Hm2DBD-NRA1                    LHFLDLFVQMFQLDTG-
Rs2DBD-NRA1                    LHFLDLFVQMFQLDTG-
Hd2DBD-NRA1                    LHFPDLFVQMFQLDTG-
Sp2DBD-NRA1                    LHFPDLYIQMFRLDDST
Sme2DBD-NRA3b                  LKFPNLYTQMFLID
Bg2DBD-NRA                     LHIPQLYAEMYNSVTTG
Ac2DBD-NRA                     LAVPQLYAEMYAS
Pc2DBD-NRA                     LEVCALYKEMFF-
My2DBD-NRA                     LTLPQMYKEMFGEKVLE
Mm2DBD-NRA                     IKVPQLFHELIIESIKE
Ga2DBD-NRA                     LQVNILFKEIFAV
Cg2DBD-NRA                     LEISPLMREVYHLNPK
Cv2DBD-NRA                     LEISPLMREVYHLNPK
Of2DBD-NRA                     LPLPQLFTEMFLND
Bf2DBD-NRA                     LKIPQLFSELHKV
Bb2DBD-NRA                     LKIPQLFSELHKV
2DBD-NRB
Ap2DBD-NRB                     VTLPALFAEINL
Pmi2DBD-NRB                    VTLPALFAEINL
Aru2DBD-NRB                    AVLPALFAEINL
Lv2DBD-NRB                     FSMPALFAEINLSS
Spur2DBD-NRB                   VSMPALFAEINLS
Cg2D2-NRB                      QEMHQLFDKLDLENPMS
Cv2D2-NRB                      QEVHQLFDKLDLENPLS
2DBD-NRC
Cr2DBD-NRC4                     LFKKNCFLMFLLRHVL
Cr2DBD-NRC1                     LFKRSSFLMFILRNIT
Cr2DBD-NRC2                     LFKRSSFLMFILRNIT
Cr2DBD-NRC3                     LFKRSSFLMFVLRNIT
NRs from Subfamily 1
hTRa (NP_955366.1)             ELFPPLFLEVFEDQEV
hTRb (NP_000452.2)             ELLPPLFLEVFED
xTRb (NP_001090533.1)          ELFPPLFLEVFED
mTRb (NP_033406.1)             ELFPPLFLEVFED
hRARa (NP_000955.1)            GSMPPLIQEMLENSEGL
mRARb (NP_035373.1)            GSMPPLIQEMLENSEGH
hRARb (NP_000956.2)            GSMPPLIQEMMENSEGH
hRARg (NP_000957.1)            GPMPPLIREMLENPEMF
```

```
mRARg (NP_001398643.1)         GPMPPLIREMLENPEMF
hPPARa (AAA36468.1)            -ALHPLLQEIYRDMY
mPPARa (CAA40856.1)            -ALHPLLQEIYRDMY
hPPARd (NP_001165289.1)        -SLHPLLQEIYKDMY
mPPARd (AAA19972.1)            -LLHPLLQEIYKDMY
mPPARg (AAA62110.1)            -SLHPLFQEIYKDLY
hPPARg (BAA18949.1)            -SLHPLLQEIYKDLY
dE78 (AAF69494.1)              --LPPLFAEIFDIPKADDEL
hRORa (NP_599023.1)            LHFPPLYKELFTSEFEPAMQIDG
mmRORa (AAB46801)              LHFPPLYKELFTSEFEPAMQIDG
mcRORa (NP_001276845.1)        LHFPPLYKELFTSEFEPAMQIDG
hRORb (CAA69929.1)             TLFPPLYKELFNPDCATACK
mRORg (NP_035411.2)            AAFPPLYKELFSTDVESPEGLSK
hRORg (AAA64751.1)             AAFPPLYKELFSTETESPVGCPS
dHR3 (NP_001246236.1)          VVFPALYKELFSIDSQQDLT
bECR (NP_001166846.1)          --LPPFLEEIWDVAEVATTH--P
dECR (NP_001163061.1)          --LPKFLEEIWDVHAIPPSVQSH
mLXRb  (NP_001272446.1)        --LPPLLSEIWDVHE
hLXRb (BAH02288.1)             --LPPLLSEIWDVHE
mLXRa (NP_001171201.1)         --LPPLLSEIWDVHE
hLXRA (AAA85856.1)             --LPPLLSEIWDVHE
hFXRa (NP_005114.1)            --FTPLLCEIWDVQ
mFXRa  (NP_033134.2)           --FTPLLCEIWDVQ
hVDR (XP_024304946.1)          MKLTPLVLEVFGNEIS
mVDR (BAA06737.1)              MKLTPLVLEVFGNEIS
hPXR (BAH02292.1)              -FATPLMQELFGITGS
mPXR (NP_035066.1)             -FATPLMQELFSSTDG
hCAR (NP_001070948.1)          -AMMPLLQEICS
mCAR (AAC53349.1)              -AMTPLLGEICS
```
